# Alleloscope: Integrative single cell analysis of allele-specific copy number alterations and chromatin accessibility in cancer

**DOI:** 10.1101/2020.10.23.349407

**Authors:** Chi-Yun Wu, Billy T. Lau, Heon Seok Kim, Anuja Sathe, Susan M. Grimes, Hanlee P. Ji, Nancy R. Zhang

## Abstract

Cancer progression is driven by both somatic copy number aberrations (CNAs) and chromatin remodeling, yet little is known about the interplay between these two classes of events in shaping the clonal diversity of cancers. We present Alleloscope, a method for allele-specific copy number estimation that can be applied to single cell DNA and ATAC sequencing data, either separately or in combination. This approach allows for integrative multi-omic analysis of allele-specific copy number and chromatin accessibility on the same cell. On scDNA-seq data from gastric, colorectal, and breast cancer samples, with extensive validation using matched linked-read sequencing, Alleloscope finds pervasive occurrence of highly complex, multi-allelic copy number aberrations, where cells that carry varying allelic configurations adding to the same total copy number co-evolve within a tumor. The contributions of such allele-specific events to intratumor heterogeneity have been under-reported and under-studied due to the lack of methods for their detection. On scATAC-seq from two basal cell carcinoma samples and a gastric cancer cell line, Alleloscope detects multi-allelic copy number events and copy neutral loss-of-heterozygosity, enabling the dissection of the contributions of chromosomal instability and chromatin remodeling in tumor evolution.

## Introduction

Cancer is a disease caused by genetic alterations and epigenetic modifications which, in combination, shape the dysregulated transcriptional programming of tumor cells^1, 2^. These somatic genomic events lead to a diverse cellular population from which clones with advantageous alterations proliferate and eventually metastasize^3^. The comprehensive study of cancer requires the integrative profiling of genetic and epigenetic changes at the resolution of single cells. We combined the analysis of two such genomic dimensions – DNA copy number and chromatin accessibility – through massively parallel single cell sequencing assays.

First, consider copy number aberrations (CNAs), through which we have derived much of our current understanding of the relationship between genome instability and tumor evolution^4^. Total copy number profiling, which estimates the sum of the copy numbers of the two homologous chromosomes, is inadequate to characterize some types of cancer genomic aberrations. Such events include the pervasive copy-neutral losses of heterozygosity (LOH)^5-8^ and the intriguing “mirrored events”^9, 10^, where cells carrying amplification of one haplotype are intermingled with cells carrying amplification of the other haplotype. While the importance of allele-specific copy number profiling has been emphasized in bulk DNA sequencing analysis^5-8, 11^, most single-cell studies profile only total copy number due to the difficulty of reliable allele-specific estimation at low per-cell coverage^12-19^. Recently, a major advance was made by Zaccaria and Raphael in the development of CHISEL^10^, a method for single-cell allele-specific copy number analysis. By default, CHISEL relies on externally phased haplotypes derived from available reference cohorts for accurate estimation of allele-specific CNAs, and found evidence for the existence of mirrored amplifications in a deeply sequenced breast tumor. This was the first time that allele-specificity of CNA events were reported at single-cell resolution, but the requirements for high sequencing depth and external phasing limit the general applicability of CHISEL. Thus, despite these recent advances, the prevalence of mirrored amplification events and the general complexity of CNA regions at single-cell allele-specific resolution are still largely unexplored.

Epigenetic modifications are also an important genomic feature of cancer. One type of epigenetic profiling is the measurement of chromatin openness, for which a variety of methods have been developed^20^ including transposase-accessible chromatin sequencing (ATAC-seq)^21, 22^, which can be performed at the bulk-tissue or single-cell level. Analysis of chromatin structure has shown that epigenetic remodeling modulates the plasticity of cells in cancer^23-27^, leads to stem-like properties^28-30^ and generates therapeutic resistance^31-34^. Since copy number alterations involve large gains and losses of chromatin, we expect the chromatin accessibility of a region to be influenced by the changes in underlying copy number. Current scATAC-seq studies estimate total copy number profiles by smoothing the read coverage and comparing position-wise against a control cell population^24, 26^. Yet this appropriate control is difficult to identify, as even between normal cells of distinct lineages, broad chromatin remodeling can lead to chromosome-level shifts in coverage (examples of this will be given in Results). There is yet no method for reliable total or allele-specific copy number profiling in scATAC-seq data, and thus, the estimation of CNA and chromatin accessibility are confounded in current analysis pipelines.

Addressing these challenges, we present Alleloscope, a method for <u>allel</u>e-<u>s</u>pecific <u>cop</u>y number <u>e</u>stimation and multiomic profiling in single cells. Alleloscope does not rely on external phasing information, and can be applied to low coverage scDNA-seq data or to scATAC-seq data with sample-matched bulk DNA sequencing data. To investigate the single cell landscape of allele-specific CNA, we first apply Alleloscope on scDNAseq data from four gastric cancer samples, four colorectal cancer samples, and two datasets from different regions of a breast cancer sample^10, 12, 35^. For five of the gastrointestinal cancer samples, results are extensively validated by 10x linked-read sequencing which provides accurate phasing information^36-38^. In these datasets, Alleloscope accurately identifies LOH and mirrored-subclonal amplification events, and furthermore, finds pervasive occurrence of highly complex multi-allelic loci where cells that carry varying allelic configurations adding to the same total copy number co-evolve within a tumor. The ubiquity of such events in all three cancer types analyzed reveal that they may be an important overlooked source of intratumor genetic heterogeneity.

Having characterized the complexity of allele-specific CNA events at single cell resolution, we turn to scATAC-seq data from two basal cell carcinoma samples with paired bulk whole exome sequencing data^26^ and a complex polyclonal gastric cancer cell line that we also analyzed by scDNA-seq. In these samples, we evaluate the accuracy of Alleloscope in genotyping and clone assignment for scATAC-seq data and demonstrate its application to the integrative analysis of CNA and chromatin accessibility.

## Results

### Overview of Alleloscope model and algorithm

First, we briefly overview Alleloscope’s method for allele-specific copy number estimation (Fig. 1). Clone assignment and integration with peak signals in scATAC-seq data will be described later. Alleloscope relies on two types of data features: coverage, derived from all reads that map to a given region, and allelic imbalance, derived from allele-informative reads that cover heterozygous loci in the region. We start with some essential definitions. For a given single nucleotide polymorphism (SNP) site, we refer to its mean coverage across cells as *bulk coverage* and its mean variant allele frequency (VAF = ratio of alternative allele read count to total read count) across cells as its *bulk VAF*. Between the two parental haplotypes, we define the term “major haplotype” as the haplotype with higher mean count across cells. Note that a haplotype may be the “major haplotype” of a sample, but have lesser copy number than the other haplotype within some cells. For each cell *i*, in any given CNA region *r*, we define two key parameters: (1) the major haplotype proportion (*θ*_*ir*_), defined as the count of the copies of the major haplotype divided by the total copy number for the region in cell *i* and (2) total copy fold change (*ρ*_*ir*_), defined as the ratio of the total copy number of the region in cell *i* relative to that in normal cells.

**Fig 1:**
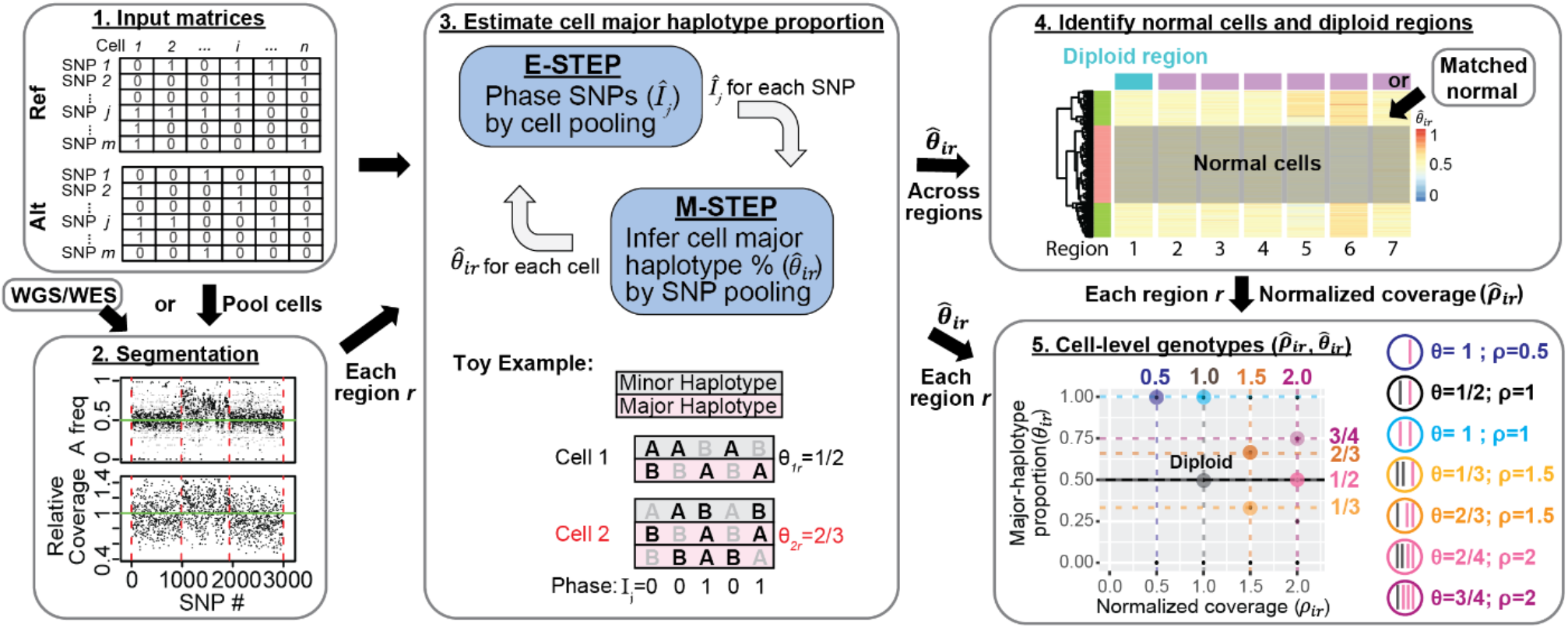
Overview of allele-specific copy number estimation of single cells with Alleloscope. 1. The algorithm operates on raw read count matrices for reference allele (Ref) and alternative allele (Alt) computed from single cell DNA or ATAC sequencing. 2. First, we obtain a segmentation of the genome based on sample-matched whole genome or whole exome sequencing data using FALCON^5^. If scDNA-seq is available, cells can be pooled to derive a pseudo-bulk. 3. We first define 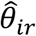 as the major haplotype proportion for cell *i* for region *r* derived from the segmentation. Then for each region *r*, Alleloscope simultaneously phase SNPs (*Î*_*j*_) and estimate cell major haplotype proportion 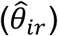 by expectation maximization (EM) algorithm. In the E-step, information is pooled across cells to estimate the phasing of each SNP. In the M-step, information is pooled across all SNPs in the region are pooled to estimate the major haplotype proportion 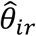 for each cell. The toy example shows a scenario with two cells for a region containing 5 SNPs, with cell 2 carrying an amplification of the major haplotype (in pink). For each cell and each SNP, alleles that are observed in a sequenced read are bolded in black (we assume that only one read is observed, reflecting the sparsity of the data). The true phase (*I*_*j*_) of the SNPs and the true major haplotype proportion (*θ*_*ir*_) are shown. 4. After 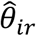 are estimated for each region *r*, 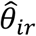 ‘s are pooled across all regions to identify candidate normal cells and candidate normal regions for computing a normalized coverage 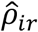 for region *r* in cell *i*. Matched normal can be specified if available. 5. Alleloscope assigns integer allele-specific copy numbers to each cell for each region based on the 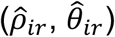 pairs.

The algorithm starts by segmenting the genome into regions of homogeneous allele-specific copy number using both the bulk coverage and bulk VAF profiles (Fig.1 Step 2). This can be achieved by multiple existing algorithms, which may be combined to increase detection sensitivity, see Methods for details. In our analyses of scATAC-seq data, the segmentation was performed on the matched bulk or single cell DNA-seq data, which ensures that the CNA regions passed to the characterization step are supported by evidence from DNA and thus are not simply due to the broad chromatin remodeling that occur in cancer.

Now consider each putative CNA region, indexed by *r*. An expectation-maximization (EM) based algorithm is used to iteratively phase each SNP and estimate the major haplotype proportion (*θ*_*ir*_) for each cell (Fig. 1, step 3). For each SNP *j* lying in the region, let *I*_*j*_ ∈ {0,1} be the indicator of whether the reference allele of SNP *j* is carried by the major haplotype (please see the toy example in Fig. 1). An initial estimate 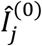 is first derived from the bulk VAF profile. Then, in iteration *t*, Alleloscope computes 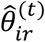 by pooling counts across sites within the region *r*, weighted by the current phasing 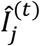, then updates the estimate of *I*_*j*_ based on 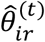 by pooling counts across cells. The estimates of *θ*_*ir*_ and *I*_*j*_ usually converge within a few iterations as described in the Methods. If matched scDNA-seq data are available for a sample sequenced by scATAC-seq, *I*_*j*_ values can be estimated from scDNA-seq and then used to compute *θ*_*ir*_ for each cell in the scATAC-seq data, enabling more information sharing between the two data types.

The estimated major haplotype proportions 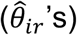, along with a preliminarily normalized coverage statistic (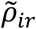 ’s, see Methods for details), are then used to identify a set of normal cells and diploid regions (Fig. 1, Step 4). This information is then used to estimate an improved relative coverage fold-change 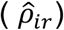 for each cell within each CNA region. If cell *i*’s true allele-specific copy numbers are homogeneous within the given region *r*, then its true value of (*θ*_*ir*_, *ρ*_*ir*_) should belong to a set of canonical points displayed in Step 5 of Fig. 1. Thus, the estimated values 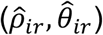 are clustered across cells and associated with one of the canonical values to yield the cell-level haplotype profiles for the CNA region. These cell-and region-specific haplotype profiles serve as the basis for clone assignment and subsequent integration with peak signals in scATAC-seq data. (Fig. 4b).

**Fig 4:**
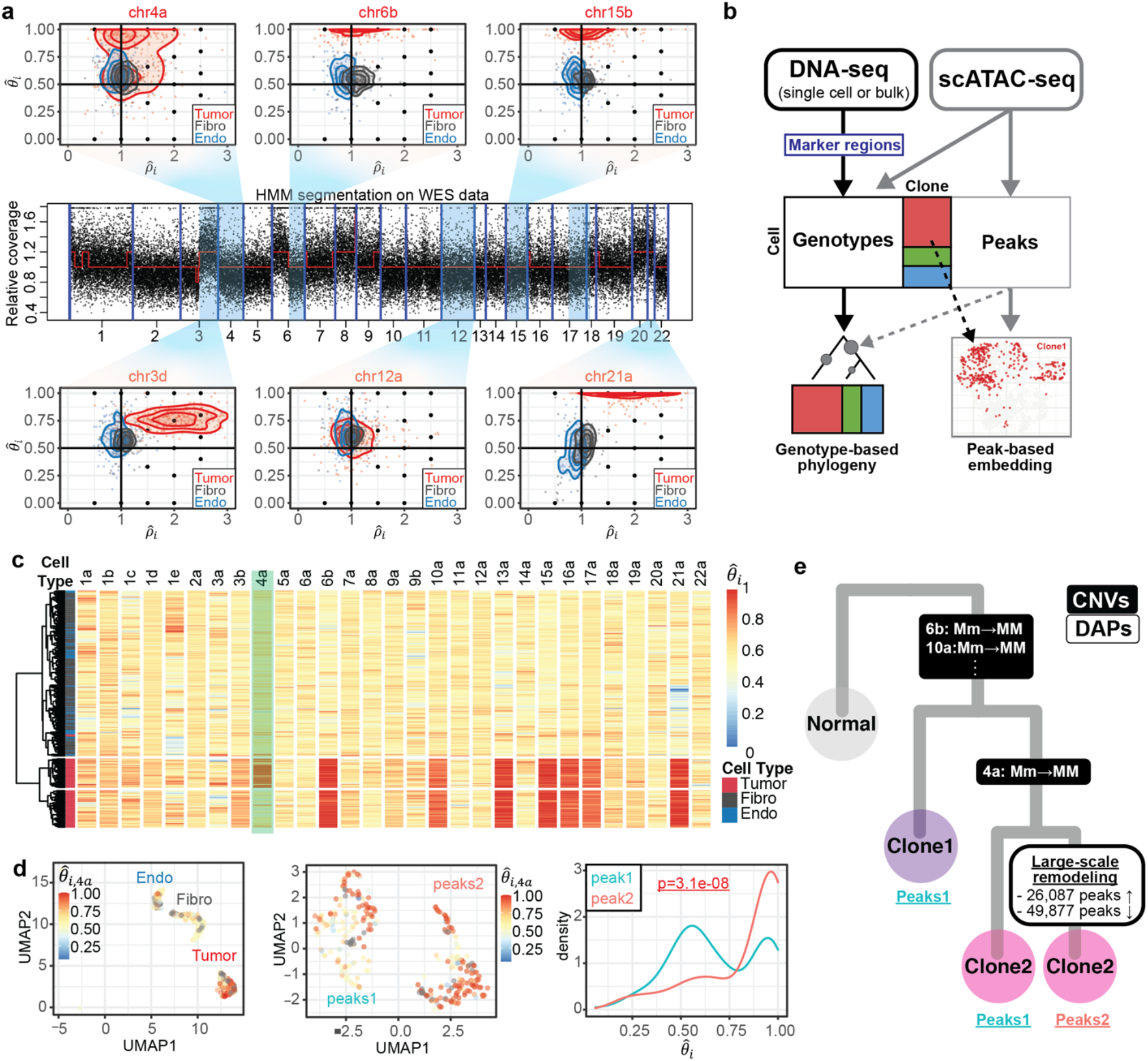
Alleloscope multiomic analysis of scATAC-seq data of a basal cell carcinoma sample (SU008^26^). (a) Genotype profiles for six example regions for cells in scATAC-seq data. The regions are taken from segmentation of matched whole exome sequencing (WES) data. Each dot represents a cell-specific 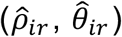 pair. Cells are colored by annotation derived from peak signals^26^, Tumor: tumor cells, Fibro: fibroblasts, Endo: Endothelial cells]. Density contours are computed for each cell type (tumor, fibroblasts, endothelial) separately and shown by color on the plot. The lower-case letters following the chromosome number in the titles denote the ordered genomic segments. (b) Pipeline for multi-omics analysis integrating allele-specific copy number estimates and chromatin accessibility peak signals on ATAC-seq data. (c) Hierarchical clustering of cells by major haplotype proportion 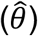 allows the separation of tumor cells from normal cells, as well as the differentiation of a subclone within the tumor cells. The marker region on chr4a separating the two tumor subclones is highlighted. (d) Integrated visualization of chr4a major haplotype proportion 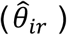 and genome-wide peak profile. Left: UMAP projection of the 788 cells in the dataset by their genome-wide peak profile, colored by 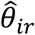. The cell type annotation (endothelial, fibroblasts, and tumor cells) is labeled in the plot. Middle: UMAP projection of only the 308 tumor cells by their genome-wide peak profile shows two well-separated clusters: peaks1 and peaks2. Right: Density of 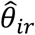 values for the peaks1 and peaks2 subpopulations. (e) Intratumor heterogeneity of SU008 is shaped by a subclonal LOH of chr4a followed by subsequent genome-wide chromatin remodeling leading to three subpopulations: Clone 1 which does not carry the chr4a LOH (peaks cluster 1), Clone 2 carrying the chr4a LOH (peaks cluster 1), and remodeled clone 2 (peaks cluster 2).

As we will show next in benchmark experiments, the above method allows accurate CNA state characterization in a wide range of scenarios. However, there is one scenario that is extremely difficult: balanced mirrored subclones at small clonal frequencies. To increase sensitivity for such events, Alleloscope allows an additional refinement step that starts with the 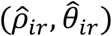 estimates and uses thresholding on 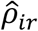 to improve the estimation of 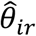. Please see Supplementary Methods for details.

### Whole genome haplotypes validate Alleloscope in scDNA-seq allele-specific copy number estimation

First, we explore the single cell landscape of allele-specific CNAs using scDNA-seq data. The scDNA-seq samples included in this study are given in Supplementary Table 1. To benchmark the phasing and allele-specific copy number state estimation accuracy, we performed matched linked-read whole-genome sequencing data on five gastrointestinal tumor samples representing different levels of chromosomal instability: P5931, P6335, *P61*9*8, P5*9*15*, and P6461. Linked-read sequencing, in which one derives reads from individual high molecular weight DNA molecules, provides variants that can be phased into extended haplotypes covering megabases in length^36-38^. To evaluate the accuracy of phasing, we compared the haplotypes estimated by Alleloscope to the haplotypes obtained from linked-read WGS. Additionally, we used the linked-read WGS haplotype with Alleloscope to derive gold-standard allele-specific copy number state assignments for each cell, which are used to assess the impact of phasing errors on accuracy in allele-specific copy number state estimation (Fig. 2a). This allows us to compute the sensitivity (recall rate of CNA regions detected in gold-standard) and specificity of a method, as defined in Supplementary Methods.

**Fig 2:**
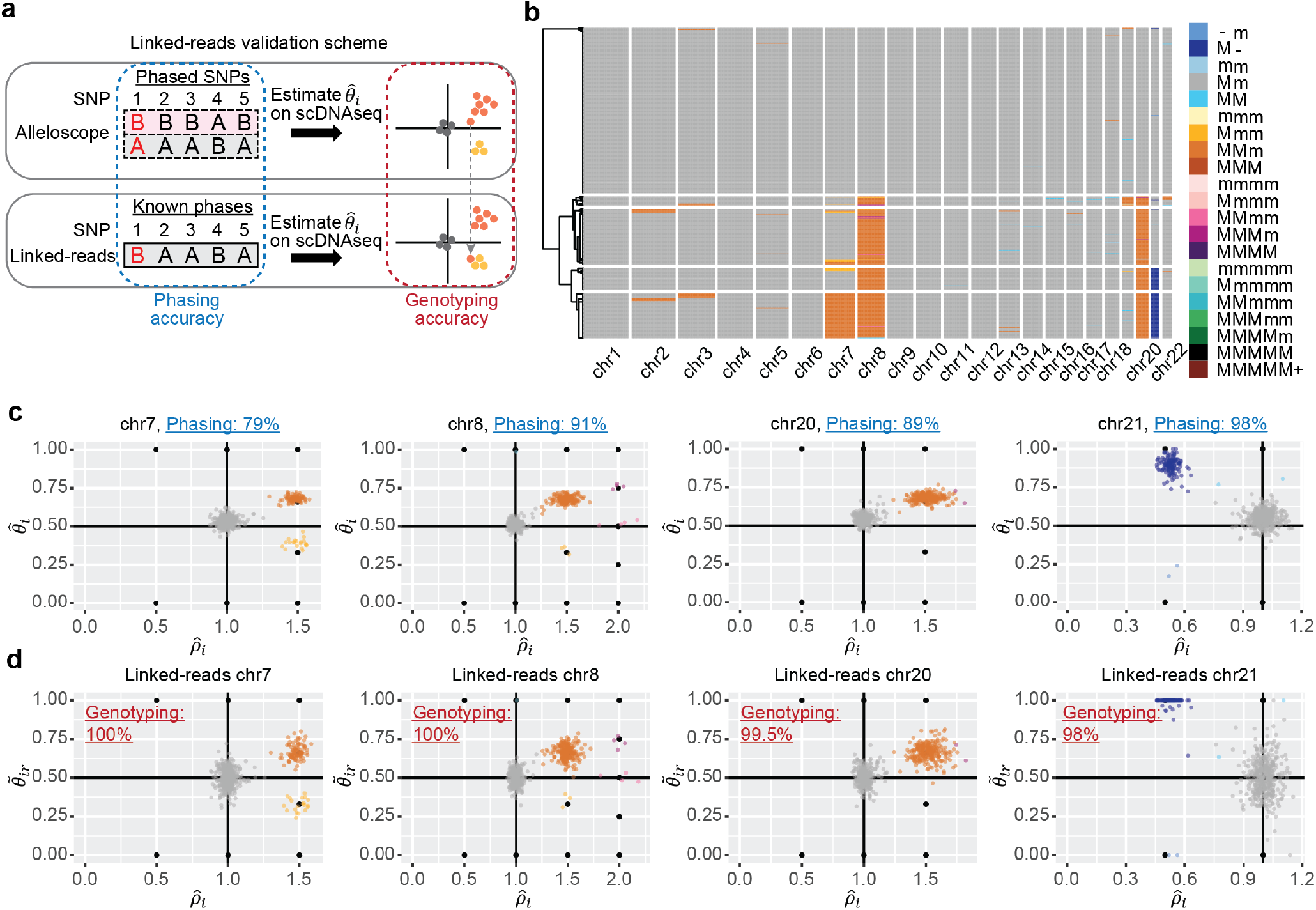
Validation of the Alleloscope results on the P5931 gastric cancer patient sample and linked-read sequencing data. (a) Illustration of the validation scheme using linked-read sequencing data. Phasing accuracy and accuracy in allele-specific copy number state estimation are used to access performance of the method. (b) Hierarchical clustering of cells in the P5931t sample based on allele-specific copy numbers given by Alleloscope, showing normal cells and 4 main clones, as well as a number of small clones marked by highly confident low-frequency mutations. M: Major haplotype, m: minor haplotype. (c) 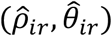 estimated by Alleloscope for four regions, colored by the inferred haplotype profile. Note that clusters fall on canonical points corresponding to discrete allele-specific copy number configurations. Phasing accuracy for each region is shown in the plot title. In the color legend, M and m represent the “Major haplotype” and “minor haplotype” respectively. (d) Similar to (c), with 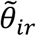 estimated using known SNP phases from matched linked-read sequencing data, colored by the haplotype profiles assigned in (c) using Alleloscope without the given phasing information. Accuracy in allele-specific copy number state estimation (Genotyping accuracy) is labeled in the plots.

We first use the gastric cancer sample P5931 to demonstrate Alleloscope and illustrate this benchmark design. The heatmap in Fig. 2b gives a bird’s-eye-view of Alleloscope’s allele-specific copy number estimates for this sample, showing it to have a relatively simple genome with few CNAs, which is expected given its MSI subtype. However, detailed inspection of the four chromosomes carrying clear CNA events, chr7, chr8, chr20, and chr21, reveals complexity at the allelic level. For each event, the scatter plots of 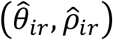 estimated by Alleloscope and colored by haplotype-specific CNA state assignment are shown in Fig. 2c. Note that the 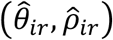 clusters fall almost directly on top of the expected canonical values (e.g. (1/2, 1) for diploid, (2/3, 1.5) for 1 copy gain of major haplotype). Interestingly, chromosomes 7, 8, and 21 each show subclonal clusters with differing allelic ratios adding up to the same total copy number, which would not have been detectable without allele-specific estimation. We denote the major haplotype of a region by “M”, and the minor haplotype by “m”. The chromosome 7 amplification exhibits two tumor subclones with mirrored-subclonal CNAs (MMm and Mmm), each subclone amplifying a different haplotype. Such a mirrored-subclonal CNA configuration is also observed for the deletion on chromosome 21 (M- and m-). The chromosome 8 amplification exhibits four tumor subclones differentiated by their haplotype profiles— MMm, Mmm, MMmm, and MMMm.

Comparing against the whole genome haplotypes derived from linked-reads, the phasing accuracy is 98% for the chr21 deletion, ∼90% for the two clonal amplifications (on chr8 and chr20), and 79% for the subclonal chr7 amplification (shown in the titles of the scatter plots of Fig. 2c). Fig. 2d shows scatterplots of 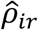 against major haplotype proportion computed using haplotypes derived from linked-read sequencing 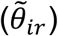, with cells colored by their state assignment given by Alleloscope without using the haplotype information (i.e. each cell has the same color as in Fig. 2c). Comparing the scatterplots in Fig. 2d to their counterparts in Fig. 2c reveals that Alleloscope’s estimated cell haplotype profiles are highly concordant with those derived directly with the haplotypes from linked-read WGS. Specifically, the concordance is ∼100% across all four events (the concordance for each event is labeled in the scatter plots of Fig. 2d). This shows that the allele-specific copy number estimation by Alleloscope is robust to errors in phasing (e.g. for chr7). We also used two extreme cases—chr2 and chr3 of P5931 to demonstrate the refinement step (Supplementary Fig. 1). For these two regions, linked-read sequencing revealed a mirrored amplification event distinguishing two subclones each comprised of <10 out of 700 cells. The differing major haplotype proportions of these two subclones, at 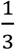 and 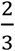, was missed by the original estimation, but accurately detected with the refinement step of Alleloscope.

So far, CHISEL is the only other comparable method for allele-specific copy number estimation with scDNA-seq data. Thus, we benchmarked Alleloscope against CHISEL by comparing each method’s estimates under default settings to the same method’s estimates using the true haplotype phasing information provided by linked-read WGS. Thus, each method’s performance is assessed using its own “gold standard” computed with linked-read WGS haplotypes. We verified that CHISEL’s gold standard is highly similar to Alleloscope’s gold standard, and thus, with the help of linked-read WGS haplotypes both methods can achieve accurate estimation that is close to the truth. The sensitivity and specificity of each method on the five samples are given in Table 1. For CHISEL, there is an extra “correction step” where the inferred clones are used to generate consensus allele-specific copy number profiles for all cells within each clone. We show CHISEL’s results with and without this correction step, as for most samples, it increases sensitivity (but decreases specificity), but for (P5931) it makes the results worse. The results obtained using different external phasing datasets and different block sizes for CHISEL also suggest that CHISEL’s accuracy depends on choices for these inputs (Supplementary Table 2). Detailed heatmaps comparing Alleloscope and CHISEL using P5931 as an example are given in Supplementary Fig. 2-3, with more discussion given in Supplementary Results. Overall, Alleloscope maintains high accuracy (both sensitivity and specificity) across the samples analyzed, substantially improving upon the current state-of-the-art.

**Table 1.** Comparison between Alleloscope and CHISEL using two benchmark strategies.

| Dataset | Method | Sensitivity | Specificity |
| --- | --- | --- | --- |
| **Benchmark with matched linked-read sequencing data** |  |  |  |
| P5915 | Alleloscope | 0.9373 | 0.9915 |
|  | CHISEL<br>(before correction) | 0.7284 | 0.9757 |
|  | CHISEL<br>(after correction) | 0.7786 | 0.9722 |
| P5931 | Alleloscope | 0.9402 | 0.9986 |
|  | CHISEL<br>(before correction) | 0.7520 | 0.9434 |
|  | CHISEL<br>(after correction) | 0.0112 | 0.9508 |
| P6198 | Alleloscope | 0.9433 | 0.9666 |
|  | CHISEL<br>(before correction) | 0.9397 | 0.9311 |
|  | CHISEL<br>(after correction) | 0.9700 | 0.9359 |
| P6335 | Alleloscope | 0.9671 | 0.9906 |
|  | CHISEL<br>(before correction) | 0.7858 | 0.9873 |
|  | CHISEL<br>(after correction) | 0.8404 | 0.9943 |
| P6461 | Alleloscope | 0.8638 | 0.9904 |
|  | CHISEL<br>(before correction) | 0.7932 | 0.9624 |
|  | CHISEL<br>(after correction) | 0.8143 | 0.9411 |
| **Benchmark with subsampling** |  |  |  |
| 50% subsample | Alleloscope | 0.9261 | 0.9905 |
|  | CHISEL<br>(before correction) | 0.9267 | 0.9383 |
|  | CHISEL<br>(after correction) | 0.9496 | 0.9446 |
| 25% subsample | Alleloscope | 0.8249 | 0.9525 |
|  | CHISEL<br>(before correction) | 0.6515 | 0.6412 |
|  | CHISEL<br>(after correction) | 0.6533 | 0.6475 |

### Further performance assessment via data downsampling and simulations

We also compared Alleloscope and CHISEL on the breast cancer sample that was analyzed in detail by Zaccaria et al^10^. This sample does not have true phasing information, but was sequenced at much higher coverage, and thus, we downsampled the reads to 50% and 25% of the original depth and compared the estimates obtained from the downsampled datasets to those obtained from the original dataset. Heatmaps of the genome-wide profiles estimated by CHISEL and Alleloscope on the original dataset, the 50% subsampled dataset and the 25% subsampled dataset are shown in Supplementary Fig. 4. For CHISEL, the clone-corrected outputs are shown since they are more accurate than the uncorrected ones. The accuracy of Alleloscope and CHISEL are shown in Table 1. We see that at 25% of the original depth, Alleloscope has a specificity of 82% as compared to CHISEL’s 65%, and sensitivity of 95% as compared to CHISEL’s 64%.

Finally, we used simulations to investigate the performance of Alleloscope over a grid of experimental parameters (number of cells, total per cell coverage, and total coverage at heterozygous SNP sites, Supplementary Fig. 5). Across experimental design settings, Alleloscope attains higher accuracy for deletions than amplifications, which is expected due to the larger change in both coverage ratio and major haplotype proportion 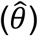 that accompanies deletions. Phasing for deletions (and other LOH events) is accurate across the board, but for amplifications it is low at 10% and 5% clonal frequencies. However, the CNA state assignment accuracy is, to a certain extent, robust to phasing error, increasing steadily with coverage, number of heterozygous SNPs, and number of cells.

### Alleloscope finds pervasive occurrence of polyclonal CNA regions differentiated by haplotype ratios

We start by considering copy neutral LOH events, which are well-known drivers of cancer evolution that can only be identified through allele-specific copy number analysis. We use the colorectal adenocarcinoma sample P6198 as illustration. The copy number heatmap in cellranger (Fig. 3a) reveals that most of the genome is amplified for this sample, but with few regions of normal copy number. One of these regions, chromosome 5, is shown by cellranger to be copy neutral for its entire length. The bulk VAF clearly separates this chromosome into two main regions, a normal region followed by a copy-neutral LOH (Fig. 3b). Concordantly, Alleloscope reveals a cluster centered at (*ρ, θ*) *=* (1,1) corresponding to copy-neutral LOH only for the region on the right (Fig. 3b). Similarly, alleloscope also revealed copy neutral LOH events in this sample on chr11 and chr16. Using haplotypes from matched linked-read WGS, all of these LOH events are validated and the per-cell LOH state assignment accuracy of Alleloscope is nearly 100% (Supplementary Fig. 6).

**Fig 3:**
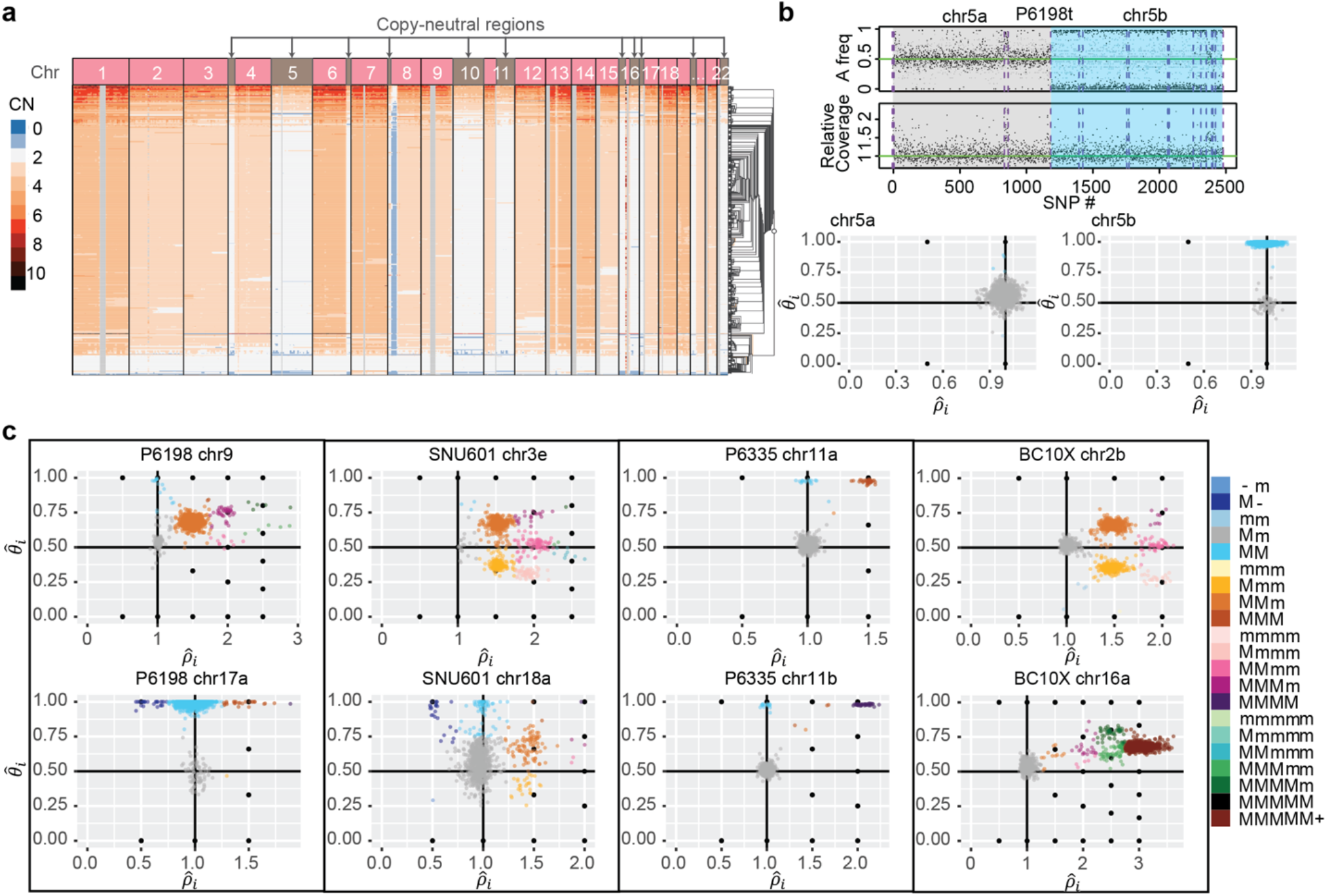
Across multiple cancer types, Alleloscope detects loss-of-heterozygosity events and multi-allelic copy number aberrations, delineating complex subclonal structure which are invisible to total copy number analysis. (a) The Cell Ranger hierarchical clustering result for P6198t with copy-neutral regions labeled (total 512 cells). (b) Top: FALCON segmentation of P6198 chr5 into two regions with different allele-specific copy number profiles. Bottom: Detailed haplotype profiles of the two regions from Alleloscope, showing that the first region is diploid across cells and the second region has a loss-of-heterozygosity for a subpopulation of cells. The a and b following the chromosome number denote two ordered segments. (c) Single cell allele-specific estimates 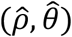, colored by assigned haplotype profiles, for select regions in the samples *P61*9*8t* (metastasized colorectal cancer sample), SNU601 (gastric cancer cell line), P6335 (colorectal cancer sample), and BC10X (breast cancer cell line). In the color legend, M and m represent the “Major haplotype” and “minor haplotype” respectively. The lower-case letters following the chromosome number in the titles denote the ordered genomic segments.

Next, we survey the overall landscape of single cell allele-specific copy number profiles among the samples analyzed. Detailed segmentation plots and heatmaps for all samples, listed in Supplementary Table 1, are in Supplementary Fig. 7-13. Across most of the samples, we observe a high prevalence of complex subclonal CNAs indicated by multiple clusters differentiated by allelic ratios for the same genomic region. Prototypical examples from P6198, SNU601, P6335 and BC10X are shown in Fig. 3c. In some regions, such as chromosome 9 of SNU601, 3q of SNU601, 2q and 16p of BC10x, we see as many as seven subclonal clusters. In many cases there are multiple clusters corresponding to the same total copy number but varying in allelic dosage. Minor subclones carrying deletion of one haplotype can be easily masked by dominant subclones carrying amplifications of the other haplotype in a bulk analysis or a single cell analysis that relies on total coverage. Overall, the high subclonal diversity in these genomic regions reveal an aspect of intratumor heterogeneity that was previously underreported.

Recurrent chromosomal instability events, affecting both haplotypes and producing gradients in haplotype dosage, is a common theme across all samples analyzed. Consider, for example, the region on chromosome 9 of P6198 (Fig. 3c), which reveals 7 subpopulations of cells: besides the normal cell cluster and the dominant tumor cell cluster with the haplotype profile MMm, there is a small cluster of cells with copy neutral LOH, two small subclones at four chromosome copies and two more at five chromosome copies. This produces major haplotype ratios of 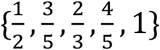 in different cells. Another example of such complexity is chromosome 3q of SNU 601 and chromosome 2q of BC10x, which share a similar pattern: two mirrored-subclonal CNAs (MMm, mmM) at total copy number of 3, as well as mirrored-subclonal CNAs (MMMm, MMmm, mMMM) at total copy number of 4, producing a gradient of haplotype ratios 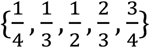. Interrogating the evolutionary route by which such diversity was achieved, Alleloscope reveals that a whole genome doubling event is likely to have taken place early in the development of BC10x and P6198, but not in the development of SNU601 (see Supplementary Fig. 4&6). Thus, the subclones at 2q in BC10x and 3q in SNU601 must have evolved through different evolutionary routes: In BC10x, the early whole-genome doubling produced the cluster MMmm, from which the other clusters of different haplotype profiles were most likely derived through successive loss and gene conversion events. On the contrary, the clusters on 3q of SNU601 were most likely a result of successive amplification events starting from the normal haplotype profile Mm. The observation of such similar clonal configurations in allele-specific copy number, in two samples coming from different cancer types (breast and gastric), suggests that convergent evolution or similar evolutionary pressures, where such haplotype dosage gradients serve as an important substrate for selection in tumor evolution, may be at play. Large studies involving more samples would be needed to characterize the evolutionary significance of such complex clonal patterns.

### Juxtaposition of single cell copy number and chromatin remodeling events by integrative scATAC-seq analysis

To demonstrate the integrative analysis of scATAC-seq data with Alleloscope, we first consider two basal cell carcinoma samples with matched whole-exome sequencing (WES) data^39^. Using the matched WES data, the genome of each sample was first segmented into regions of homogeneous bulk copy number (Fig. 4a, middle panel shows the segmentation for SU008). Alleloscope was then applied to the scATAC-seq data to derive allele-specific copy number estimates of each cell in each region. Scatterplots of 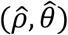 for five example CNA regions and one control region (chr12) from SU008 are shown in Fig. 4a. For this sample, peak profiles characterizing chromatin accessibility separated the cells confidently into three main clusters: 308 tumor cells, 259 fibroblasts and 218 endothelial cells with the cell type identity retrieved from the original study. Since the normal fibroblast and epithelial cells are not expected to carry broad copy number events, they provide a “null scenario” for comparison to the tumor cells. Density contours for each cell type are shown in the 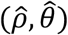- scatterplots (Fig. 4a). The 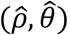 values clearly separate the tumor cells from the normal cells for each CNA region, with the tumor cell cluster positioned at canonical points in each case, indicating that these statistics used by Alleloscope can accurately distinguish amplifications and loss-of-heterozygosity events in scATAC-seq data. In particular, Alleloscope differentiated the cells that carry the putative copy neutral LOH events in regions 4a, 6b, and 15b through shifts in major haplotype proportion. Note that normal cells, which are not expected to carry broad chromosome-scale CNAs, exhibit chromosome-level deviations in total coverage due to broad chromatin remodeling but with no significant difference in their 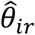 distribution, as exemplified by the chr6b region. The coverage in large genomic bins across the genome was further investigated for the two normal cell types in scATAC-seq data (Supplementary Fig. 14; Supplementary Methods). Comparing one normal cell type against the other, many regions exhibit broad shifts in relative coverage. Thus, relying solely on shifts in coverage, without complementary shifts in major haplotype proportion, would lead to false positive copy number detections for scATAC-seq data.

By assigning allele-specific CNA profiles to single cells in scATAC-seq data, Alleloscope allows the integrative analysis of chromosomal instability and chromatin remodeling as follows (Fig. 4b): The scATAC-seq data, paired with bulk or single-cell DNA sequencing data, allows us to detect subclones. In parallel, a peak-by-cell matrix can be computed following standard pipelines. Then, the subclone memberships or CNA profiles can be visualized on the low-dimensional embedding of the peak matrix, and the subclones can be further compared in terms of peak or transcription factor motif enrichment. Precise haplotype profiles for each subclone then allow us to identify significantly enriched/depleted peaks after accounting for copy number differences, thus delineating events that are uniquely attributable to chromatin remodeling.

Hierarchical clustering using major haplotype proportion 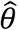 identifies the tumor cells from the normal cells for both SU006 (Supplementary Fig. 15) and SU008, and clearly delineates a subclone in SU008 marked by a copy-neutral LOH event on chr4a (Fig. 4c). Focusing on SU008, we call the cell lineage that carries the chr4a LOH event clone-2, and the other lineage clone-1. In parallel, clustering by peaks cleanly separates the tumor cells from the epithelial cells and fibroblasts (Fig. 4d: left), and further, demarcates two distinct clusters in the tumor cells (peaks-1 and peaks-2) (Fig. 4d: middle). What is the relationship between the peaks-1 and peaks-2 clusters obtained from peak signals to the two clones delineated by chr4a LOH? Coloring by chr4a major haplotype proportion 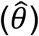 on the peaks-derived UMAP shows that the LOH in this region is carried by almost all of the cells in peaks-2 but only a subset of the cells in peaks-1 (Fig. 4d: middle). This can also be clearly seen in the density of 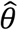 (Fig. 4d: right): While 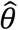 is heavily concentrated near 1 for peaks-2, it is bimodal for peaks-1. Since clone-1 and clone-2 are differentiated by a copy-neutral event, this separation by peaks into two clusters is not driven by broad differences in total copy number. Since clone-2 is split into two groups of distinct peak signals, we infer that the chromatin remodeling underlying the divergence of the peaks-2 cells is likely to have occurred in the clone-2 lineage, after the chr4a LOH event (Fig. 4e). In this way, Alleloscope analysis of this scATAC-seq data set allowed us to overlay two subpopulations defined by peak signals with two subpopulations defined by a subclonal copy-neutral LOH, and infer their potential temporal order.

### Integrative analysis of clonal evolution and altered chromatin accessibility for a complex polyclonal gastric cancer cell line

The gastric cancer cell line SNU601 exhibits complex subclonal structure, as evidenced by multiple multiallelic CNA regions (chr3e and chr18a are shown in Fig. 3c). In addition to scDNA-seq, we also performed scATAC-seq on this sample to profile the chromatin accessibility of 3,515 cells at mean coverage of 73,845 fragments per cell. This allows us to compare the allele-specific copy number profiles obtained by scATAC-seq with those given by scDNA-seq and integrate the two data types in a multi-omic characterization of this complex tumor.

First, we segmented the genome and estimated the allele-specific copy number profiles of single cells at each segment for both the scATAC-seq and scDNA-seq data, following the procedure in Fig. 1 with some modifications due to the lack of normal cells to use as control for this sample (see methods). Fig. 5a shows the relative total coverage, pooled across cells from scDNA-seq. Fig. 5b shows 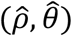-scatterplots for five example CNA regions in scDNA-seq and scATAC-seq. Compared to the scATAC-seq data, the scDNA-seq data has about 8-fold higher total read coverage and 7-fold higher heterozygous site coverage per cell. Thus, while subclones corresponding to distinct haplotype profiles are cleanly separated in the scDNA-seq data, they are much more diffuse in the scATAC-seq data. Yet, cluster positions in scATAC-seq roughly match those in scDNA-seq. As expected, the 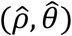-scatterplots reveal the high level of chromosomal instability in this sample, with each region exhibiting multiple clusters of different haplotype structures that indicate the existence of subclones carrying mirrored events and, for some regions, the variation of haplotype dosage over a gradient across cells.

**Fig 5:**
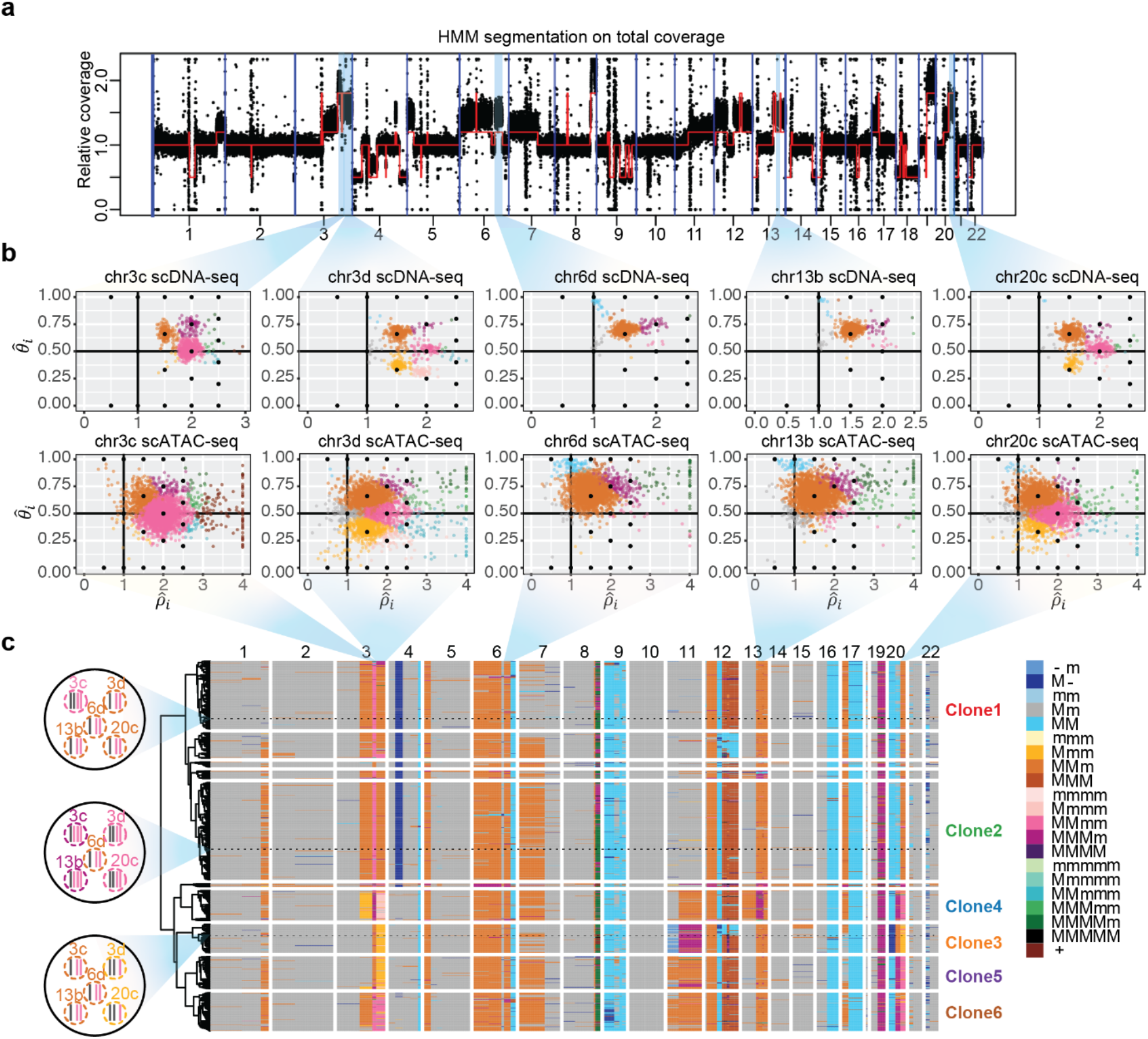
Alleloscope analysis of scDNA-seq and scATAC-seq data reveals complex subclonal heterogeneity in the SNU601 gastric cancer cell line. (a) Genome segmentation using HMM on the pooled total coverage profile computed from scDNA-seq data. (b) Single cell allele-specific copy number profiles 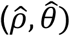 for five regions in scDNA-seq and scATAC-seq data. Cells are colored by haplotype profiles according to legend in Fig. 5c. (c) Tumor subclones revealed by hierarchical clustering of allele-specific copy number profiles from the scDNA-seq data. Genotypes of the five regions shown in Fig. 5b, for three example cells, are shown in the left. The haplotype structures for the 5 regions in Fig. 5b of three cells randomly chosen from Clone 1, 2, and 3, are shown to the left of the heatmap. In the color legend, M and m represent the “Major haplotype” and “minor haplotype” respectively. The six clones selected for downstream analysis in scATAC-seq data are labeled in the plot.

Fig. 5c shows the hierarchical clustering of cells from scDNA-seq based on their allele-specific copy number profiles, revealing the subclonal structure and the co-segregating CNA events that mark each subclone. For each cell in each region, Alleloscope also produces a confidence score for its assignment to different haplotype profiles (Supplementary Fig. 16). Based on visual examination of the confidence scores at the marker regions in both the ATAC and DNA sequencing data sets, we identified 6 subclones for further investigation (Clones 1-6 labeled at the right of the heatmap). The allele-specific copy number profiles allow us to manually reconstruct the probable evolutionary tree relating these 6 clones under the following three rules:

1. Parsimony: The tree with the least number of copy number events is preferred.
2. Monotonicity: For a multi-allelic region with escalating amplifications (e.g. Mm, MMm, MMMm), the haplotype structures were produced in a monotonic order (e.g. Mm→ MMm→ MMMm) unless a genome doubling event occurred.
3. Irreversibility of LOH: Once a cell completely loses an allele (i.e. copy number of that allele becomes 0), it can no longer gain it back.

The evolutionary tree, thus derived, is shown in Fig. 6b. The mirrored-subclonal amplifications on chr3q, the deletion on chr4p, and the multiallelic amplification on chr20q allowed us to infer the early separation of clones 3-6 from clones 1-2. Subclones 3-6 are confidently delineated by further amplifications on chr3q, chr20q, chr11, chr13, and chr17. Note that high chromosomal instability led to concurrent gains of 1q and 7p in both the Clone 1-2 and Clone 3-6 lineages. We also observed a large number of low-frequency but high-confidence CNA events indicating that ongoing chromosomal instability in this population is spawning new sporadic subclones that have not had the chance to expand.

**Fig 6:**
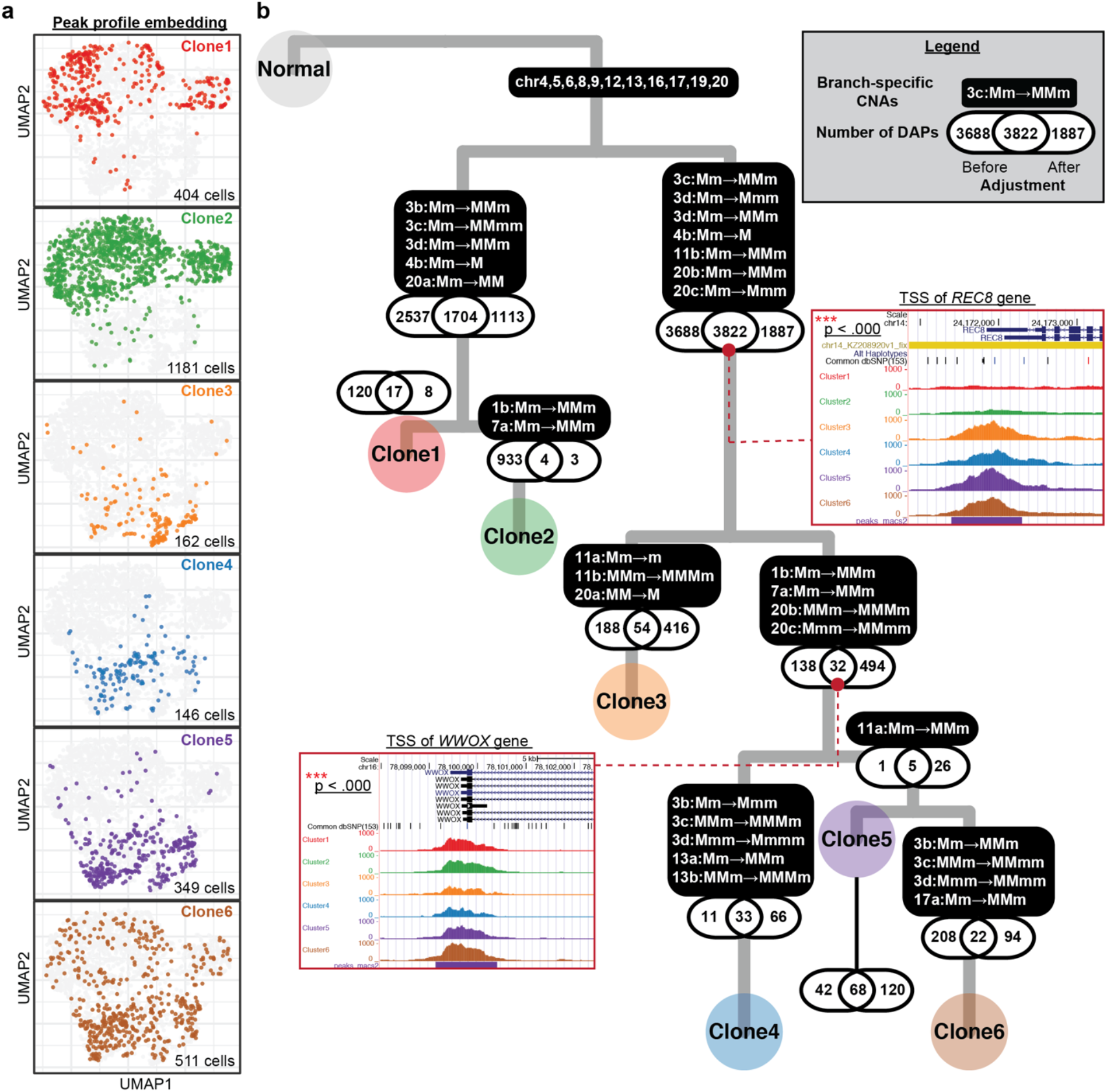
Integrative analysis of allele-specific copy number and chromatin accessibility for SNU601 ATAC sequencing data. (a) UMAP projection of genome-wide scATAC-seq peak profile on 2,753 cells. The same group of cells were clustered into one of the six subclones based on their allele-specific copy number profiles across the 10 selected regions. Cells in different subclones are labeled with different colors, using the same color scheme as that for the subclone labels in Fig. 4c. The number of cells colored in each UMAP is shown at the bottom-right corners. (b) A highly probable lineage history of SNU601, with copy number alterations (CNAs) and differentially accessibility peaks (DAPs) marked along each branch. P-values of the tests for association between DAPs and CNAs are shown along each branch. For two example DAP genes, pooled peak signals for each subclone are shown as inset plots.

We now turn to scATAC-seq data, focusing on the 10 marker regions which, together, distinguish Clones 1-6: chr1b, 3b-d, 4b, 7a, 11b, 13b, and 20b-c. The 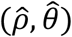 values computed by Alleloscope allows us to directly assign allele-specific copy number profiles to each cell for each region, as well as subclone labels to each cell, with posterior confidence score. The subclone assignment utilizes a Bayesian mixture model that pools information across the 10 marker regions. Despite the low accuracy in per-region genotyping, when information is pooled across the 10 marker regions, 81.6% of the 2,753 cells after filtering can be assigned to a subclone with >95% posterior confidence (Supplementary Fig. 17, the number of ATAC cells confidently assigned to each clone are shown in Fig. 6a.). These subclone assignments for each cell, and cell-level haplotype profiles for each region, can now be integrated with peak-level signals.

Following the scheme in Fig. 4b, we computed the Uniform Manifold Approximation and Projection (UMAP) coordinates for the scATAC-seq cells based on their peak profiles, which enables us to visualize the difference in chromatin accessibility profiles among individual cells (Fig. 6a). UMAP scatterplots colored by clone assignment show that the 6 clones exhibit marked differences in their chromatin accessibility profiles (Fig. 6a): While Clone 1 and Clone 2 are concentrated at the top half of the UMAP, Clones 3-5 are positioned almost exclusively at the bottom half. Clone 6, which exhibits more variance, is also significantly enriched at the bottom half of the UMAP. Among Clones 3-5, Clone 3 has a distinct chromatin accessibility profile that is mostly concentrated at the bottom tip, Clone 4 is positioned higher, while Clone 5 contains cells that are similar to both clones 3 and 4. We expect some of these peak-level differences to be driven by CNAs.

To delineate the peaks that differ between clones, and to distinguish peak differences that are not accountable by CNAs, we developed a statistical test to adjust for underlying copy numbers to identify differentially accessibility peaks (DAPs) between the cell populations on the two branches (Fig. 6b). The test is based on a generalized log-likelihood ratio test with specific adjustment terms for the copy number profile of each of the two subclones (Methods). The same test, but neglecting coverage adjustment, was also performed for comparison. Along each branch, three numbers are shown for DAPs— 1. DAPs identified only before adjustment; 2. DAPs identified both before and after adjustment; 3. DAPs identified only after adjustment (Fig. 6b). For the smaller subclones (Clone 3,4,5), low coverage limits the detection power and thus limits the DAP counts in both categories. Yet, juxtaposing DAP and CNA events along the tumor phylogeny yields insights: Along most lineages, adjustment for CNA-induced broad shifts in coverage removes a large number of peaks whose differential-accessibility signal is explainable by the underlying change in copy numbers. Interestingly, CNA adjustment also identifies new DAPs whose signals were obscured by a change in copy number in the opposite direction. This argues for the importance of CNAs as a mechanism underlying subclonal differences in chromatin accessibility in this tumor.

Nevertheless, along some branches we find a large number of DAPs remaining after adjustment (i.e. DAPs identified both before and after adjustment), and thus must be due to other mechanisms. Two example DAPs of this latter category are shown as insets in Fig. 6b, with full list given in Supplementary Table 3. The first example is a peak at the transcription start site (TSS) of the REC8 gene, which is located on chr14 where no apparent CNAs were observed across the six major subclones. The TSS of REC8 is open in clones 3-6 but closed in clones 1-2 (p-value<0.0001). REC8 is a gene encoding a meiosis-specific cohesion component that is normally suppressed in mitotic proliferation, and its role in cancer has recently gained increasing attention and controversy: While Yu et al.^40^ found the expression of this gene to suppress tumorigenicity in a gastric cancer cell line, McFarlane et al.^41^ postulated that it may be broadly activated in some cancers where it generates LOH by reductional segregation. The opening of the TSS of REC8, stably maintained in Clones 3-6, suggests that meiotic processes may underlie the increased chromosomal instability of this multiclonal lineage. The second example is a peak at the TSS of the WWOX gene, located on chr16, which is significantly depleted in Clone 3 (p-value<0.0001). Although chr16 has LOH across all tumor cells, there are no detectable subclonal differences, and thus we don’t expect the decrease in accessibility at WWOX for subclone 3 to be due to a large copy number event. Since WWOX is a well-known tumor suppressor whose down-regulation is associated with more advanced tumors^42, 43^, its decrease in accessibility suggests a more aggressive phenotype for Clone 3. Overall, these two examples show how Alleloscope can be used to dissect the roles of CNA and chromatin-level changes in the identification of gene targets for follow-up study.

## Discussion

Despite recent advances in the application of single cell sequencing to cancer, we are still far from understanding the diversity of genomes that are undergoing selection at the single cell level. Notably, little is yet known about the intratumor diversity of allelic configurations within CNA regions, and to what extent the diversity of cells in chromatin accessibility can be attributed to diversity in allele-specific copy number. Here, we presented Alleloscope, a new method for allele-specific copy number estimation that can be applied to single cell DNA and ATAC sequencing data (separately or in combination). Through a combination of matched linked-reads whole genome sequencing, downsampling-based benchmark experiments, and simulations, we comprehensively assessed the accuracy of Alleloscope and benchmarked against CHISEL, which is currently the only other method for scDNA-seq allele-specific copy number profiling. Detailed discussion about the two methods can be found in Supplementary Results.

We applied Alleloscope to a panel of breast, colorectal, and gastric tumors and cancer cell lines, where it revealed an unprecedented level of allelic heterogeneity within hypermutable CNA regions. In these regions, subclones reside on a gradient of allelic ratios that is unobservable in total copy number analysis. In simple cases, these hypermutable regions contain mirrored subclones, as previously identified^9, 10^, but are often much more complex. We observed multiple instances of convergent evolution involving recurrent CNA events affecting the same region, some verified by linked read sequencing. In accordance with the findings in Watkins et al.^44^, we found using Alleloscope that chromosomal instability drives the formation of subclones not only in primary tumors but also after metastasis. Alleloscope was also applied to a scDNA-seq dataset^15^ generated from a different sequencing protocol (Supplementary Fig. 18; Supplementary Methods), further demonstrating the general applicability of the approach.

Having established the allelic complexity of CNAs at single cell resolution, we next applied Alleloscope to scATAC-seq data, thus enabling the combined study of clonal evolution and chromatin accessibility. First, we considered the analysis of a public data set consisting of two basal cell carcinoma samples, for which matched bulk whole-exome sequencing data was used for initial genome segmentation upon which single cell CNA genotyping was then conducted in the scATAC-seq data. Here we showed that Alleloscope can detect amplifications, deletions, and copy-neutral LOH events accurately in scATAC-seq data, and was able to find a subclone delineated by a copy-neutral LOH event. Juxtaposing this subclone assignment with peak signals allowed us to detect a wave of genome-wide chromatin remodeling in the lineage carrying the LOH. Next, we applied Alleloscope to a complex polyclonal gastric cancer cell line with matched scDNA-seq data. By overlaying peak signals with subclones delineated by allele-specific copy number estimates, we developed a statistical test to adjust for copy numbers in identifying DAPs. Comparison between the testing results before and after copy number adjustment suggests that much of the intratumor heterogeneity in chromatin accessibility can be attributed to CNAs. Focusing on subclone-enriched peaks with copy number adjustment allowed the prioritization of genes for downstream follow-up.

Alleloscope can potentially be applied to the integration of single cell data of other modalities, for example scATAC-seq and scRNA-seq data, to investigate the relationships between clonal evolution, chromatin remodeling, and transcriptome. To facilitate experimental design for single cell omics sequencing protocols, we investigated the performance of Alleloscope under different experimental parameters (number of cells, total per cell coverage, and total coverage at heterozygous SNP sites). Coverage at heterozygous SNP sites is an especially important consideration for the design of scRNA-seq and scATAC-seq studies, for which shifts in total coverage is an unreliable proxy for underlying DNA copy number. For scATAC-seq, the lower heterozygosity within peak regions led to lower number of reads mapping to heterozygous loci as compared to scDNA-seq, and this resulted in noisier subclone detection. Most of the current scRNA-seq technologies only sequence either the 3’ or 5’ end of the mRNA transcripts, which limits the number of heterozygous SNP sites covered by reads. The latest developments in single cell long read sequencing^45-47^ and single cell multimodal sequencing^48^ herald new analysis opportunities with this method.

## Methods

### ScDNA-seq Data Sets and Pre-processing

The ten 10x scDNA-seq datasets analyzed in this study are summarized in Supplementary Table 1. P5931, P5847 and P6461 scDNA-seq data were generated using the method described in the previous study^35^. We applied the Cell Ranger DNA pipeline (https://support.10xgenomics.com/single-cell-dna/software/overview/welcome; beta version: 6002.16.0) for sample demultiplexing, read alignment, CNA calling and visualization. Most data were aligned to the GRCh38 reference genome except for the two processed BC10x datasets (GRCh37). For the tumor samples with a matched normal sample, the GATK HaplotypeCaller was used to call heterozygous SNPs on the matched normal samples. Otherwise, SNPs were retrieved on the tumor sample themselves. Next, we applied VarTrix to efficiently generate SNP-by-cell matrices for both reference and alternative alleles of the identified SNPs.

To select high-quality SNPs, we filtered out the SNPs with <5 reads for P5846 and P5847, <10 reads for P5915 and P5931, <15 reads for P6335 and P6461, <20 reads for P6198, SNU601, and ther BC10X samples based on the number of SNP detected for each sample. Additionally, SNPs located in the centromeres and telomeres were excluded. To exclude cells that might undergo apoptosis or cell cycles, noisy cells labeled by the Cell Ranger tool were excluded.

### Single-cell ATAC Data sets, Sequencing and Preprocessing

The scATAC-seq datasets analyzed in this study are summarized in Supplementary Table 4. We used the following procedure to generate the SNU601 scATAC-seq dataset.About 400,000 cells were washed with RPMI media and centrifuged (400g for 5 min at 4°C) twice. The supernatant was removed and resuspended with chilled PBS + 0.04% BSA solution. The resuspended pellet was centrifuged (400g for 5min at 4°C), and the supernatant was removed again. Then 100 µL of chilled Lysis Buffer (10 mM Tris-HCl (pH 7.4), 10 mM NaCl, 3 mM MgCl_2_, 1% BSA, 0.1% Nonidet P40 Substitute, 0.1% Tween-20 and 0.01% digitonin) was added and carefully mixed 10 times. The tube was incubated on ice for 7 min. After incubation, 1 mL of chilled Wash Buffer (10 mM Tris-HCl (pH 7.4), 10 mM NaCl, 3 mM MgCl_2_, 1% BSA and 0.1% Tween-20) was added and mixed 5 times followed by centrifugation of nuclei (500g for 5 min at 4°C). After removing the supernatant, nuclei were resuspended in chilled Nuclei Buffer (10X Genomics), filtered by Flowmi Cell Strainer (40uM) and counted using a Countess II FL Automated Cell Counter. Then the nuclei were immediately used to generate scATAC-seq library.

ScATAC-seq library was generated using the Chromium Single Cell ATAC Library & Gel Bead Kit (10X Genomics) following the manufacturer’s protocol. We targeted 3000 nuclei with 12 PCR cycles for sample index PCR. Library was checked by 2% E-gel (Thermofisher Scientific) and quantified using Qubit (Thermofisher Scientific). Sequencing was performed on Illumina NextSeq500 using NextSeq 500/550 High Output Kit v2.5 (Illumina).

Raw sequencing reads of the SNU601 scATAC-seq sample was de-multiplexed with the 10x Genomics Cell Ranger ATAC Software (v.1.2.0; https://support.10xgenomics.com/single-cell-atac/software/pipelines/latest/algorithms/overview) and aligned to the GRCh38 reference genome. The aligned scATAC-seq data of the two pre-treatment basal cell carcinoma samples (BCC;SU006 and SU008) were retrieved from GSE129785^26^. To obtain all potential SNPs for the SU006 and SU008 samples, GATK Mutect2 was used to call single-nucleotide variants (SNVs) on the bam files deduplicated by the Picard toolkits of both the t-cell dataset and the tumor dataset from the same tumor. All SNVs from the paired tumor-normal datasets were combined and the read counts of these SNPs were quantified for each cell in the tumor scATAC-seq dataset. For the SNU601 scATAC-seq data, we instead quantified the read counts of the two alleles of the SNPs more reliably called from the paired normal scDNA-seq data. Then the SNP-by-cell matrices for both reference and alternative alleles were generated using Vartrix for all the scATAC-seq datasets. To select potential germline SNPs, we further filtered out the SNVs <5 reads for the SU008 sample and <10 reads for the SU006 sample. SNPs with extreme VAF values <0.1 or >0.9 were also excluded for both samples. Because we used the phasing information from the paired scDNA-seq data to assist the estimation of the haplotype structures for the SNU601 scATAC-seq data, we did not filter out SNPs with extreme VAF values for the scATAC-seq data. To improve the quality of the downstream analysis for SNU601 scATAC-seq data, we filtered out cells <5 reads and the SNPs <5 reads.

### Linked-read sequencing and data processing

The five samples with the linked-read sequencing data were acquired as surgical resections following informed consent under an approved institutional review board protocol from Stanford University. Samples were subjected to mechanical and enzymatic dissociation as previously described, followed by cryopreservation of dissociated cells^33^.

Cryofrozen cells were rapidly thawed in a bead bath at 37 °C. Cell counts were obtained on a BioRad TC20 cell counter (Biorad, Hercules, CA) using 1:1 trypan blue dilution. Between 1.5-2.5 million total cells were washed twice in PBS. Centrifugation was carried out at 400g for 5 minutes. PBS was removed and cell pellets were frozen at −80°C. DNA extraction was carried out on cell pellets following thawing using either MagAttract HMW DNA Kit (P5931) or AllPrep DNA/RNA Mini Kit (Qiagen Inc., Germantown, MD, USA) as per manufacturer’s protocol. Quantification was carried out using Qubit (Thermofisher Scientific).

Sequencing libraries were prepared from DNA using Chromium Genome Reagent Kit (v2 Chemistry) (10X Genomics, Pleasanton, CA, USA) following manufacturer’s instructions. Sequencing was performed using Illumina HiSeq or NovaSeq sequencers using 150×150 bp paired end sequencing and i7 index read of 8 bp. Long Ranger (10X Genomics; v2.2.0) was used to align reads, call and phaseSNPs, indels and structural variants.

### Segmentation

The first step of Alleloscope is to segment the genome into regions with different CNA profiles. The appropriate segmentation algorithm depends on what samples are available. First, matched bulk DNA sequencing data (WGS/WES) or pseudo-bulk data from scDNA-seq data can be segmented using FACLON^5^, a segmentation method that jointly models the bulk coverage and bulk VAF profiles, if a matched normal sample is available. To accommodate segments from rare subclones, methods that integrate shared cellular breakpoints in CNA detection for scDNA-seq^49^ can improve sensitivity. If a normal sample is unavailable, Alleloscope instead uses an HMM-based segmentation method. The HMM method, which operates on the binned counts of pooled cells, assumes a Markov transition matrix on four hidden states representing deletion, copy-neutral state, single-copy amplification and double-copy amplification:

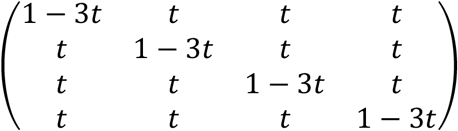, where *t* = 1 × 10^−6^ by default. The segmentation plots in the study were generated by using the HMM algorithm. With the paired sample, the P6198 tumor sample was segmented using FALCON on the SNPs >30 reads with default parameters. Besides FALCON and HMM methods, users can choose the methods that work best for specific datasets. For example, we also tried ASCAT^11^ on the P6198 sample for comparison (Supplementary Fig. 10).

### Whole-exome sequencing (WES) data processing

The WES data of the two paired tumor-normal samples (SU006 and SU008) were obtained from PRJNA533341. Raw fastq files were aligned to the GRCh37 reference genome using bwa-mem^50^ with duplicate reads removed using the Picard toolkits^51^. The copy number calls of paired normal-tumor samples were obtained using Varscan2^52^. Then the HMM algorithm was applied to segment the genome.

### SNP Phasing and Single-cell Allele Profile Estimation per region

For each region *r* after segmentation, an expectation-maximization (EM)-based method is used to iteratively phase each SNP and estimate cell-specific allele-specific copy number states for scDNA-seq and scATAC-seq datasets. “Major haplotype” is defined as the haplotype with higher aggregate copy number in the sample. Let *I*_*j*_ indicate whether the reference allele of SNP *j* is located on the major haplotype and *θ*_*ir*_denote major haplotype proportion of cell *i* for region *r*. The EM model iterates the E-step and the M-step. The complete log likelihood of the model is

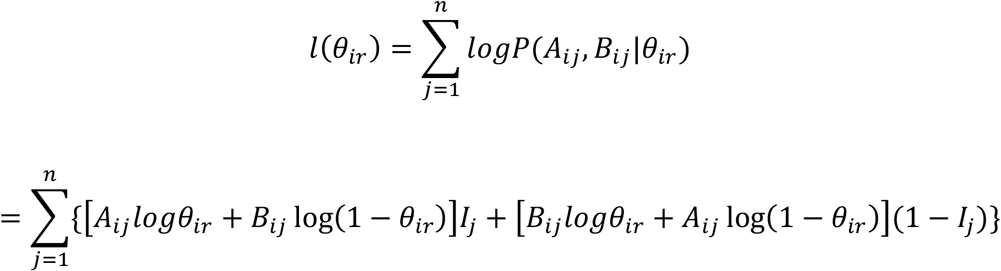

where *A*_*ij*_ and *B*_*ij*_ are the observed read counts for the reference and alternative alleles of cell *i* on SNP *j*. In the E-step, we calculate the expected value of the posterior probability of the hidden variable *I*_*j*_.

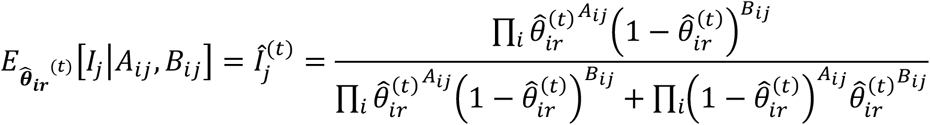

where 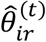 is the parameter from the t^th^ iteration. In the M-step, 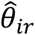 is updated by solving

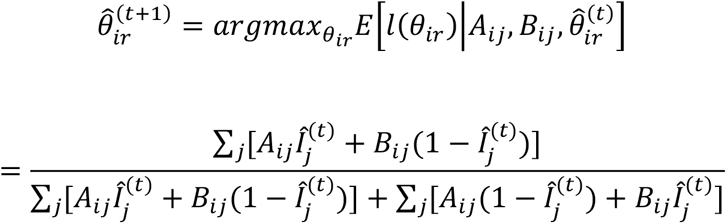

where 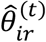 and 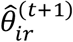 are from two successive iterations of EM. An initial estimate 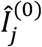 is first derived from the bulk VAF profile, and the two steps are iterated until converge. The maximum number of SNPs was set as 30,000 by default. For the SNU601 scATAC-seq dataset, the phases estimated in the paired scDNA-seq dataset (*Î*_*j*_ ’s) were directly applied to estimate the 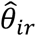 of the cells in the scATAC-seq data. To have enough power, we included the cells with ≥ 20 reads covering the identified SNPs for each region.

### Selecting normal cells and normal regions for single-cell Coverage Normalization

To compute the relative coverage change for each cell in region *r* 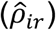, normal cells and diploid regions are required for normalization. The 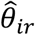 ‘s inferred from the EM-based algorithm are used to help identify normal cells and diploid regions. To identify normal cells, the cells were divided into k largest groups (*k* = 5 by default) by hierarchical clustering. The c^th^ cluster is considered normal if it has the minimum distance calculated by

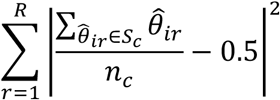

where *S*_*c*_ represents 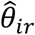 values of the cells in the c^th^ cluster, and *n*_*c*_ is total cell number in the c^th^ cluster. All cells in the c^th^ cluster are considered as candidate normal cells.

Putative diploid regions are next identified in each cluster by considering both 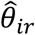 and coverage. Alleloscope computes the first measurement (*d*_*cr*_) as the sum 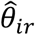 distance of the cells in the c^th^ cluster for each region *r*.

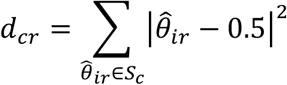

Because regions with both haplotypes equally amplified can also have small *d*_*cr*_, adjusted raw coverages (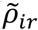; defined as below) of cell *i* and region *r* are also considered in diploid region selection.

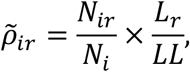

where for cell *i* and region *r, N*_*ir*_ is the total read counts and *N*_*i*_ is the total read counts across regions. *L*_*r*_ is length of the region *r* and *LL* is total length of the genome. For region *r*, cells with 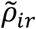 values larger than the 99^th^ percentile are assigned the 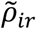 values equal to the 99^th^ percentile across the cells. The second measurement (*m*_*cr*_) used to select diploid regions in the c^th^ cluster is the mean 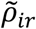 for each region *r*

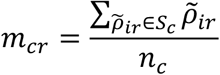

where *S*_*c*_ here represents 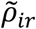 values of the cells in the c^th^ cluster. Then Alleloscope identifies potential diploid regions for each cluster by raking the sums of *d*_*cr*_ ranks and *m*_*cr*_ ranks. Excluding the c^th^ cluster identified as the normal group, Alleloscope proposed a list of candidate diploid regions across the clusters by selecting the majority region.

Because coverage on scATAC-seq data is confounded by epigenetic signals, chromosome 22 for SU008 and chromosome 18 for SU006 were directly selected as normal regions based on the matched WES data. Cells were considered as normal or tumor cells based on the epigenetic signals. For the SNU601 scATAC-seq dataset, chromosome 10 was selected as the normal region based on the paired scDNA-seq data.

### Cell-level CNA state estimation

The cell-level allele-specific copy number profiles are defined by both relative coverage change 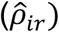 and major haplotype proportion 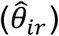 for region *r*. After the normal cells and a control region are identified, cell-specific relative coverage change in region r is calculated as

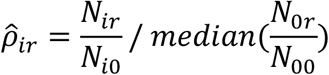

where *N*_*ir*_ is total read counts in region *r* and *N*_*i0*_ is total read counts in a reference region for cell *i. N*_*0r*_ is a vector denoting total read counts in region *r* of all identified normal cells and *N*_*00*_ is a vector denoting total read counts in the same reference region of all identified normal cells. For P6461 with no diploid region, 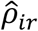 was adjusted by timing 2 for the abnormal cells. Because SNU601 is a tumor cell line with no normal cells in the dataset, *N*_*0r*_ and *N*_*00*_ were calculated from the cells in the normal P6198 sample instead for the scDNA-seq data. For SNU601 scATAC-seq data, we aligned the distribution of the 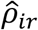 values in paired scDNA-seq data to the distribution of the 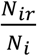 values for each region to get the normalized 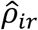 in the scATAC-seq data. The normalized 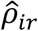 values for the scATAC-seq data were computed by

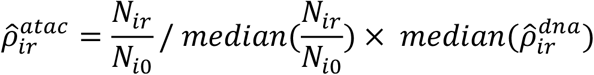

Cells with extreme 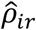 values larger than the 99^th^ percentile and smaller than the first percentile across the cells were excluded for each region. With the 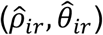 pairs, cells in the scDNA-seq data can be classified into haplotype profile *g* with the expected (*ρ*_*g*_, *θ*_*g*_) values based on minimum distance. With noisier signals in the scATAC-seq data, the haplotype structures identified in the paired scDNA-seq data can help guide the genotyping for each region. For region *r*, the posterior probability that cell *i* carries haplotype profile *g* was

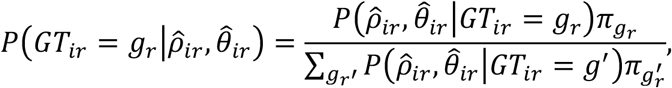

where *g*_*r*_ denotes the haplotype profile observed in region *r* in the paired scDNA-seq data and 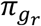 denotes the prior probability that a randomly sampled cell carrying the *g*_*r*_ haplotype profile. A uniform prior was used for 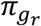. In the formula,

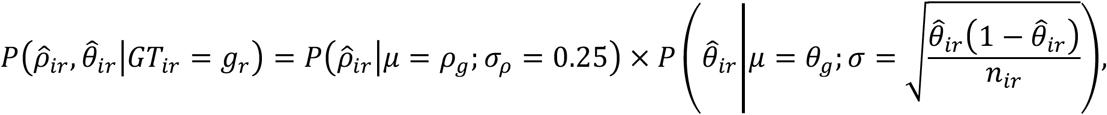

where *n*_*ir*_ is total read counts in region *r* for cell *i*. For scATAC-seq data, the haplotype profile of cell *i* in region *r* was estimated by maximizing the above posterior probability. The haplotype profiles of each region are visualized using different colors in scatter plots for both scDNA-seq and scATAC-seq data with the confidence scores calculated using the distance of the points to the canonical centers and the standard deviations.

### Validations using matched linked-read sequencing data for each region

We performed validations on five samples using matched linked-read sequencing data. Two strategies were used for the validations. First, to assess the phasing accuracy of region *r*, the largest haplotype block within region *r* in the matched linked-read sequencing data was selected for comparison, ensuring that the included SNPs are phased with respect to one another. Then the *Î*_*j*_ estimates for SNPs within region *r* in the scDNA-seq data were converted to binary indicator *Ĩ* _*j*_

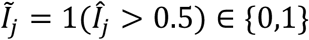

Using the major/minor haplotype setting, all SNPs in the largest haplotype block in the matched linked-read sequencing data were placed accordingly to retried *I*_*j*_ as the gold standard. Comparing the binary estimates *Ĩ* _*j*_ and *I*_*j*_, the phasing accuracy for region *r* was computed as follows:

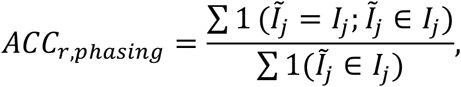

Secondly, we evaluated the accuracy for cell-level CNA state estimation for region *r*. To do this, for each cell in region *r* we compared the estimated haplotype profiles 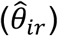 to the haplotype profiles (*θ*_*ir*_) obtained by plugging in known phases provided by the matched linked-read sequencing data as the gold standard. Similarly, the haplotype block with the largest number of SNPs was used here. The *θ*_*ir*_’s, used as the gold standard, were retrieved by directly plugging in known phases *I*_*j*_, provided by the matched linked-read sequencing data (explained in the first strategy). Then the accuracy for cell-level CNA state estimation of region *r* is computed as

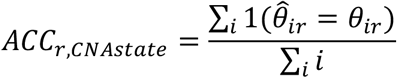

### Simulations and Power Analysis

For a simulated region, let n be the number of cells, m be the number of heterozygous SNPs, *θ* be the major haplotype proportion, and *μ*_*i*_ be the total coverage of cell i sampled from the cells on chr7 in the P5931 tumor sample. For cell i, we simulated total coverage of SNP j (*μ*_*ij*_) using a Poisson distribution

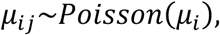

where *i* = 1∼*n*. Parallelly, phases of SNP j (*I*_*j*_) were simulated under a Bernoulli distribution

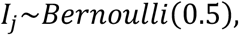

where *I*_*j*_ indicates whether a reference allele is on the major haplotype for SNP j, and j=1∼m. Using *μ*_*ij*_ and *I*_*j*_, simulated read counts of reference alleles of SNP j in cell i (*A*_*ij*_) were simulated under a Binomial distribution

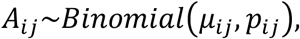

where *p*_*ij*_ is the proportion of the reference allele at loci j in cell i with the values shown in the following table

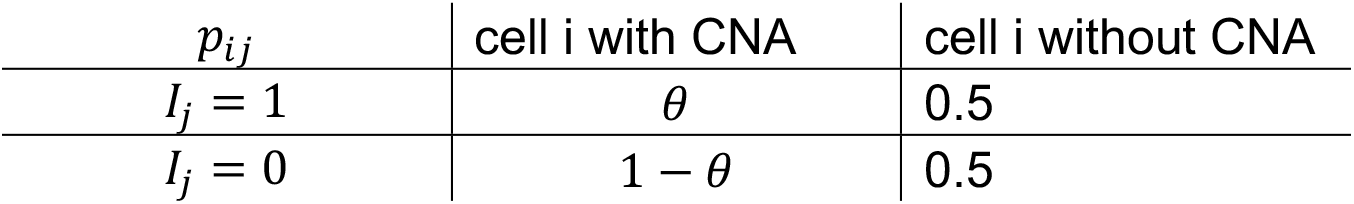

Then simulated read counts of alternative alleles of SNP j in cell i (*B*_*ij*_) were retrieved by

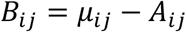

In the first simulation used to illustrate distribution of the estimates from Alleloscope, we fixed the cell number n to be 1,000, the SNP number m to be 10,000 which are typical in real datasets. *θ* was set to be 1 and 0.66 for cells carrying deletion and one-copy amplification respectively with the purity equal to 0.5. On the simulated *A*_*ij*_ and *B*_*ij*_ matrices Alleloscope estimated phases for each SNP and CNA states for each cell. Distribution of the estimated values versus the true values are visualized using boxplots.

To explore the effects of SNP numbers, cell coverage, cell numbers, and purity, power analysis was performed for one-copy deletion and one-copy amplification scenarios. We assessed the accuracy for phasing and cell-level CNA state estimation under the following scenarios: SNP numbers from 1,000 to 50,000, mean coverage from 0.01 to 0,05 for each cell, cell number from 500 to 2500. For different scenarios, we assessed the effect of three purity: 0.5, 0.1, and 0.01, reflecting from larger subclones to rare subclones. All parameters remained the same as those in the previous paragraph except for the parameters that were assessed. Phasing accuracy was calculated by comparing true *I*_*j*_ ‘s and estimated *Î* _*j*_ ‘s in the region. If *Î* _*j*_ ≥ *0*.5, the values were considered as 1. Otherwise, the values were considered 0. On the other hand, the accuracy of cell CNA state estimation was the clustering accuracy using the estimated 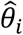 values. Cells with 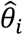 values smaller than the midpoints between true *θ* of normal cells (*θ*_o_ = *0*.5) and true *θ* of carriers (*θ*_*del*_ = 1; *θ*_*amp*_ = *0*.66) were considered as normal cells; otherwise, cells were considered as carriers. The clustering accuracy was calculated by comparing the clusters to the true cell states.

### Cell Lineage Reconstruction

For scDNA-seq data, cell-specific haplotype profiles from Alleloscope across the genome are used to reconstruct cell lineage trees. The “Gower’s distance*r* is calculated using “cluster*r* R package on the nominal haplotype profiles between cells. The ‘pheatmap’ R package was used to perform hierarchical clustering on the distance using the “ward.D2*r* method. Because fewer SNPs lead to higher variance in 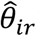 ‘s, we by default included segments with more than 2,000 SNPs identified. For visualization, each segment was plotted with its length proportional to 5,000,000 bins, and the heights of the clustering tree were log-transformed.

To explore tumor subclonal structure in the two BCC scATAC-seq datasets, we instead clustered the cells using 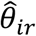 values, which are orthogonal to the CNA signals based on total coverage, across the segments with more than 500 SNPs. Then hierarchical clustering is performed on the Euclidean distance using the “ward.D2*r* method.

Because the subclones for the SNU601 sample were identified from the scDNA-seq data, we adopted a supervised strategy to assign each cell in the SNU601 scATAC-seq dataset into different subclones. First, we identified 10 marker regions-- chr1b, 3b-d, 4b, 7a, 11b, 13b, and 20b-c that help to differentiate the cells into the six major subclones based on the subclone specific copy number profiles from the scDNA-seq data. Combining the haplotype profiles across the ten regions for each cell enables assignment of the cells into one of the six subclones with high confidence. The posterior probability of cell *i* coming from clone *k* was

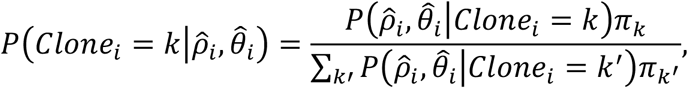

where *k* ∈ {1∼6} for the six clones and *π*_K_ is the prior probability that a randomly sampled cell coming from the *k*^*th*^ clone, which was set to uniform in the analysis.. In the formula,

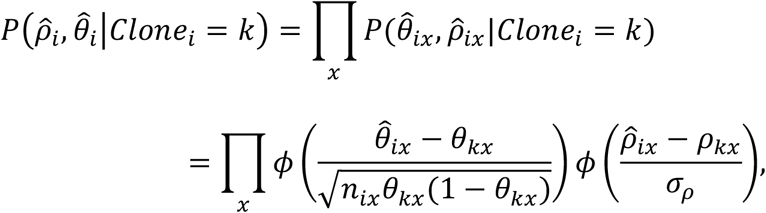

where *x* indexes the ten marker regions; 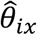 and 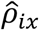 are the estimated major haplotype proportion and relative coverage for cell *i* in the scATAC-seq data; *θ*_*kx*_ and *ρ*_*kx*_ are the “known values*r* for specific haplotype profiles for clone *k* derived from the paired scDNA-seq data; and *n*_*ix*_ is the number of total read counts in the *x*^*th*^ marker region for cell *i*. Each cell was assigned into one of the six subclones by maximizing the posterior probability with the confidence score being the posterior probability of the assigned clone.

### ScATAC-seq data analysis

To investigate the relationships between allele-specific CNAs and chromatin accessibility, we processed the peak signals in addition to the allele-specific CNAs for each cell in scATAC-seq data. For the two public BCC samples, the peak-by-cell matrices were obtained from GSE129785. We log-transformed the count matrices, selected the peaks in >10% cells, regressed out cell total coverage for each peak by linear regression, and projected the cells onto the UMAP plot using genome-wide peak signals^53^. The cell type identify for each cluster was retrieved from the previous study^26^. To further explore intratumor heterogeneity, we selected the cells labeled as tumor cells, repeated the pre-processing steps described above, and projected the tumor cells onto the UMAP plot. Then DNA level information and epigenetic signals for each cell were integratively visualized.

For the SNU601 scATAC-seq dataset, scATAC-pro^54^ was used to call peaks and generate the peak-by-cell matrix. We first filtered out the cells that have proportions of fragments on the detected peaks <0.4 and or total peaks outside of the range 15,000∼100,000, and filtered out the peaks observed in less than 10% of cells. After the matrix was log-transformed, we regressed out cell total coverage for each peak by linear regression. Using Louvain clustering, the cells in two clusters with extreme total fragments were removed. After cell filtering, we repeated the process and projected the cells onto the UMAP plot. Then the clonal assignment based on the DNA information and the peak signals were integrated at the single-cell level

### Differentially accessible peaks (DAPs) identification with copy number adjustment for scATAC-seq data

To identify differentially accessible peaks (DAPs) after accounting for copy number differences between two clones, we developed a statistical test based on a generalized log-likelihood ratio (LLR) statistic. We first define some necessary terms: Given a segmentation of the genome, for a given clone *c*, let *θ*_*c*_ *=* {(*L*_*R*_, *Z*_*R*_)} be the copy number profile, which can be expressed as a vector of region lengths (*L*_*R*_) and average copy numbers (*Z*_*R*_) across all regions *R*. For peak *k*, we define function *f*_*K*_(*θ*_*c*_) to be the proportion of genomic DNA in the peak region based on the copy number profile of clone *c*, computed as follows:

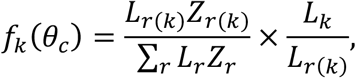

where *r*(*k*) is the region *r* to which peak *k* belongs and *L*_*”*_ is the length of peak *k*.

For peak *k* and clone *c*, let *Y*_*cK*_ be the total read count across the cells in the clone, and let *N*_*c*_ be the total read count across all peaks summed across the cells in the clone. *Y*_*cg*_ can be modeled by Binomial sampling,

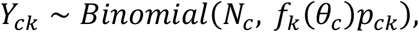

where, *p*_*ck*_ is the relative probability of detecting reads in the peak k region after adjusting for the copy numbers. Between clones with different copy number profiles, differences in *p*_*ck*_ is evidence for chromatin remodeling. Thus, pairwise comparisons were performed for each branch (clone *c*_(_ versus clone *c*_*\**_) and for each peak, using the generalized likelihood ratio test with the hypothesis 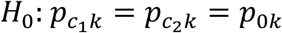.and 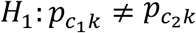.

Let *l*(*p*) denote the log-likelihood function of *Y*_*cK*_ under this model, and 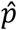 denote the MLE. The LLR for GLRT follows the Chi-squared distribution as below with derivation of MLE estimates in the Supplementary Methods.

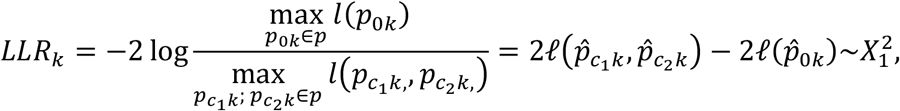

Using the GLRT, a peak is considered as a DAP if its FDR adjusted p-value<0.01 for the pairwise clonal comparison.

Simultaneously, the GLRT under the binomial model without copy number adjustment was also performed for comparison:

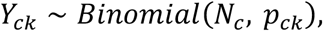

The same criterion was used to identify DAPs for each pairwise clonal comparison (FDR adjusted p-values<0.01). Each DAP was further mapped to the genes that are potentially regulated based on the *±* 2,000 bp distance on the genome.

## Supporting information

Supplementary_Alleloscope

## Data availability

The patient scDNA-seq and linked-read sequencing data generated for this study are available under dbGAP identifier phs001711. The scATAC-seq dataset is available in the National Institute of Health’s SRA repository under accession PRJNA674903. There are no restrictions on data availability or use. The other patient scDNA-seq data were obtained from dbGAP under accession phs001818.v3.p1^35^ and phs001711^12^. The cell line scDNA-seq dataset was from the Sequence Read Archive (SRA) under accession *PRJNA49880*9. The public scATAC-seq data and whole exome sequencing data were obtained from the SRA under accession PRJNA532774^26^ and PRJNA533341^39^.

## Code availability

Alleloscope is available on GitHub at https://github.com/seasoncloud/Alleloscope.

## Acknowledgements

The work is supported by the National Institutes of Health [P01HG00205ESH to B.T.L., S.M.G. and H.P.J., 5R01-HG006137-07 and 1U2CCA233285-01 to C-Y.W. and to N.R.Z., 1R35HG011292-01 to B.T.L.]. Additional support to HPJ came from the Research Scholar Grant, RSG-13-297-01-TBG from the American Cancer Society, Clayville Foundation and the Gastric Cancer Foundation.

## Author contributions

C.-Y.W. and N.R.Z. conceived the computational methods and designed the study with help from H.P.J. C.-Y.W. developed and implemented the computational methods and conducted all data analyses. B.T.L. helped in data interpretation. B.T.L., H.K. and A.S. performed all related sample preparation and sequencing. S.M.G. performed data pre-processing and coordinated data transfer. H.P.J. advised all experiments and data collection. C.-Y.W., N.R.Z, and H.P.J. wrote the manuscript. All authors read and approved the final manuscript.

