## Supplementary_Alleloscope for "Alleloscope: Integrative single cell analysis of allele-specific copy number alterations and chromatin accessibility in cancer"

### Supplementary Information

Fig. 16. Confidence scores for the genotype assignment of each cell in each region for the SNU601 scDNA-seq dataset. 20

Fig. 17. Distribution of the posterior confidence scores of subclone assignment for the 2,753 cells from SNU601 scATAC-seq. 21

Fig. 18. The segmentation plot and heatmap of the genome-wide haplotype profiles for the HM-SNS sample. 22

**Supplementary Tables 23**

Table 1. Summaries of the scDNA-seq datasets 23

Table 2. Performance of CHISEL with or without external phasing information. 23

Table 4. Summaries of the scATAC-seq datasets 24

**Supplementary Results 25**

Benchmark 1: Assessment of Alleloscope and CHISEL using whole genome haplotypes derived from linked read sequencing. 25

Benchmark 2: Assessment of method robustness by downsampling. 28

**Supplementary Methods 31**

Second-stage estimation 31

Benchmarks of Alleloscope and CHISEL 31

Assessment of coverage in large genomic bins for scATAC-seq 34

Inference of MLE estimates for the copy number adjustment model 35

HM-SNS data preprocessing and analysis 36

#### Supplementary Figures

##### Supplementary Figure 1.

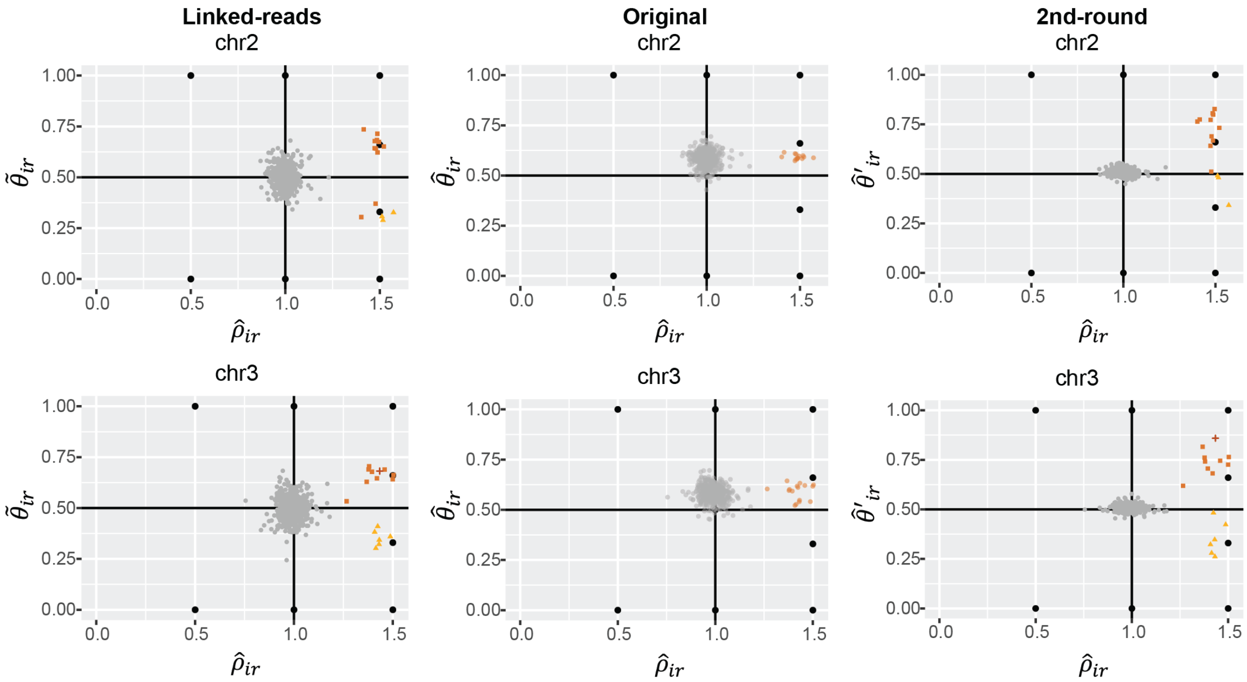

**Supplementary Fig. 1: Demonstration of Alleloscope's refinement step on two example regions of P5931.** For chr2 and chr3 (each row) of P5931, the haplotype profiles of three scenarios are shown (each column): $(\hat{\rho}_{ir}, \tilde{\theta}_{ir})$ values with $\tilde{\theta}_{ir}$ computed by plugging known SNP phases from matched linked-read sequencing data, colored by the inferred haplotype profile in c (left); $(\hat{\rho}_{ir}, \hat{\theta}_{ir})$ values with $\hat{\theta}_{ir}$ originally estimated by Alleloscope; and $(\hat{\rho}_{ir}, \hat{\theta}'_{ir})$ values with $\hat{\theta}'_{ir}$ estimated by Alleloscope's refinement step. The colors and shapes represent different haplotype profiles from the original and second-round estimation.

**Supplementary Figure2.**

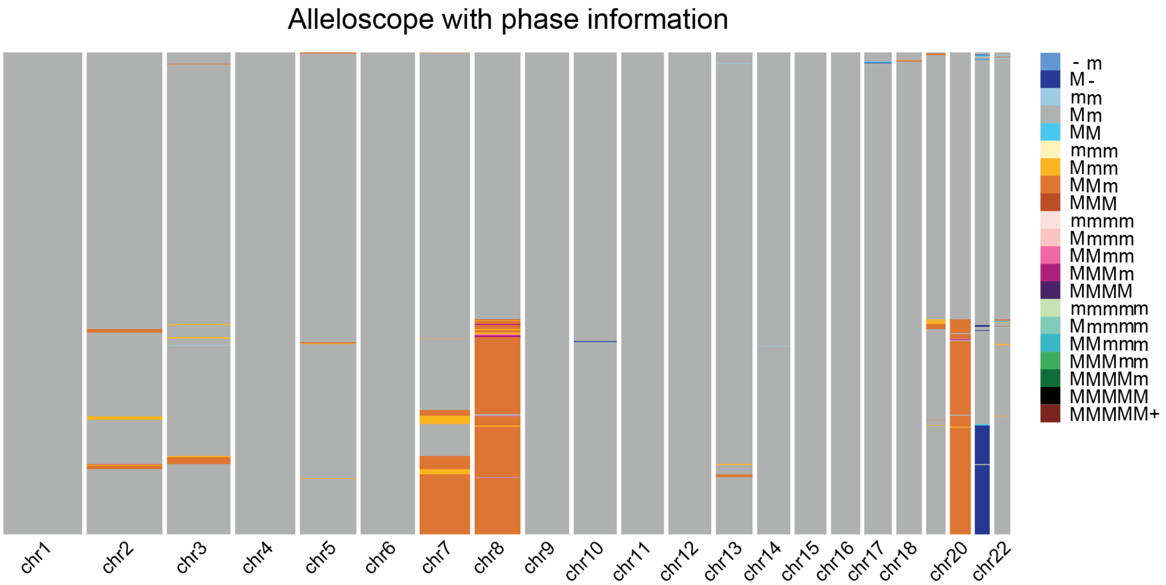

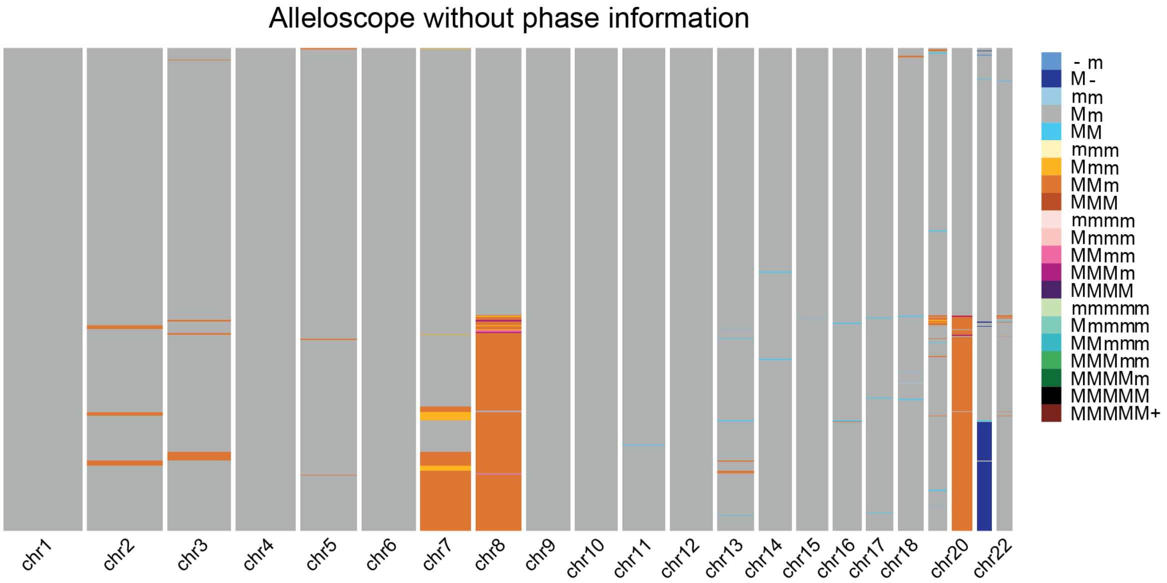

**Supplementary Fig. 2: The estimated genome-wide haplotype profiles by Alleloscope with (top) and without (bottom) known phases for the P5931 sample.** In the color legend, M and m represent the “Major haplotype” and “minor haplotype” respectively. The cells in both plots are ordered by their ordering in the linked-read-based heatmap.

#### Supplementary Figure 3

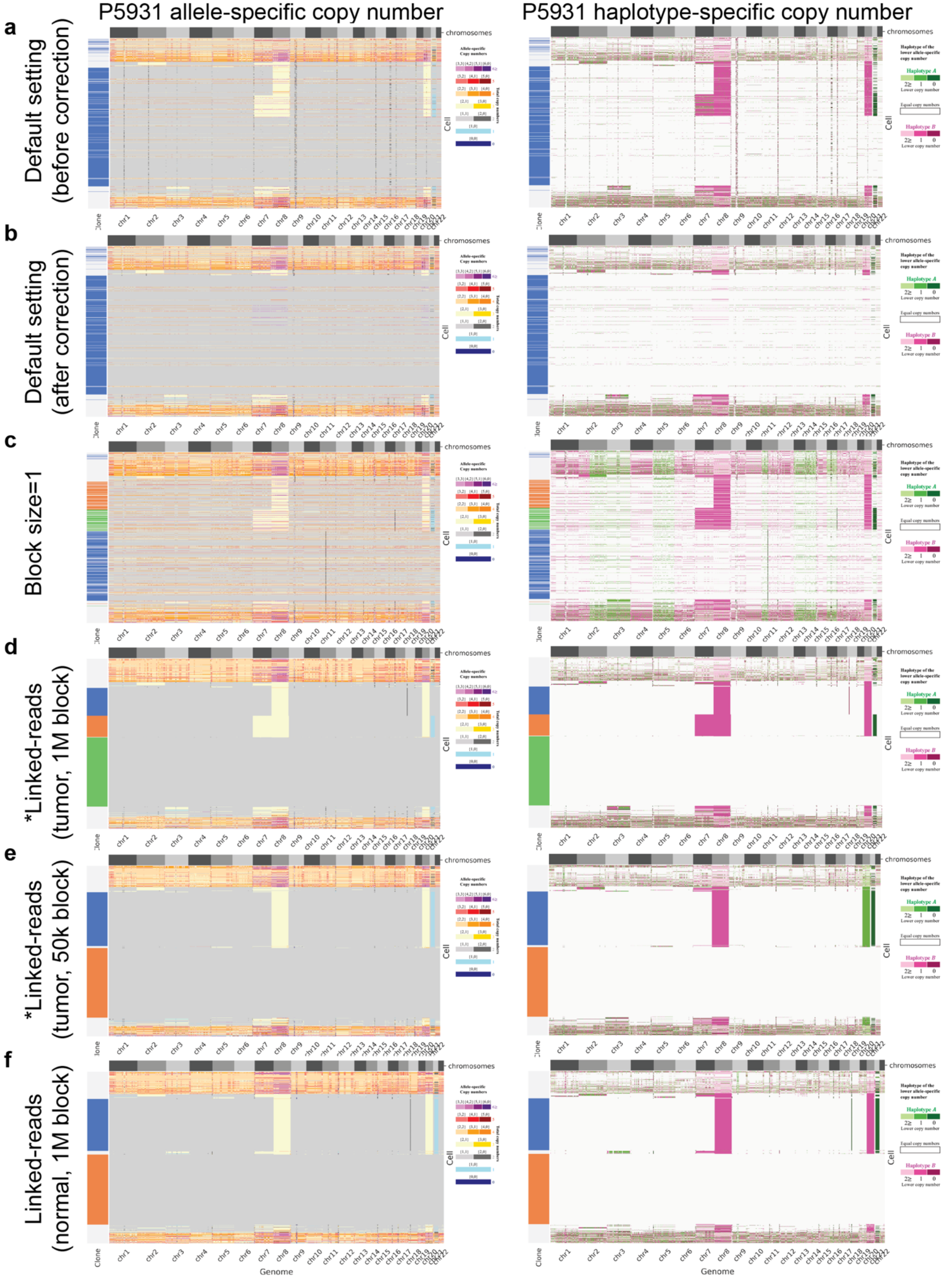

**Supplementary Fig. 3: The estimated genome-wide haplotype profiles by CHISEL with and without known phases for the P5931 sample under different settings.** For each result (each row), CHISEL gives two heatmaps: the allele-specific copy number state (left), and the major/minor haplotypes where there is an allelic imbalance (right). CHISEL includes an additional “correction step”, in which the inferred clones are used to generate consensus allele-specific copy number profiles for all cells within each clone. (a) The result before correction estimated using the default setting with the cell’s ordering from d. (b) The result after correction estimated using the default setting with the cell’s ordering from d. (c) The result after correction estimated using a block size=1 with the cell’s ordering from d. (d) The result by plugging in known phases provided by the matched tumor linked-read sequencing with 1M block size. (e) The result by plugging in known phases provided by the matched tumor linked-read sequencing with 50k block size. (f) The result by plugging in known phases provided by the matched normal linked-read sequencing with 1M block size.

**Supplementary Figure 4.**

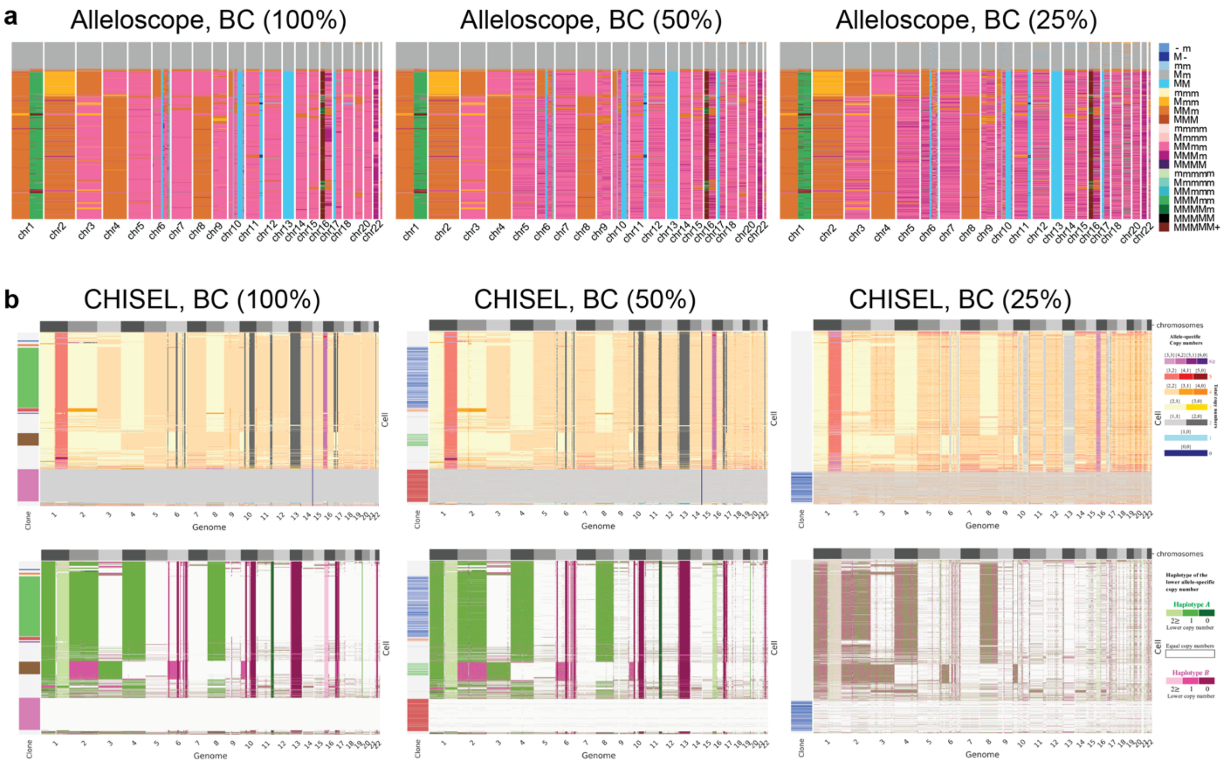

**Supplementary Fig. 4: Down-sampling results of both Alleloscope and CHISEL on a high-coverage scDNA-seq dataset of a breast cancer sample (section E).** The results on the original dataset (left), on the 50% subsampled dataset (middle) and on the 25% subsampled dataset (right), given by Alleloscope (a) and CHISEL (b).

For Alleloscope, M and m represent the “Major haplotype” and “minor haplotype” respectively in the color legend. For CHISEL, two heatmaps are given: the allele-specific copy number state (top), and the major/minor haplotypes where there is an allelic imbalance (bottom). The corrected plots are shown here. The cells in the plots of 50% and 25 subsampled datasets are ordered by their ordering in the linked-read-based heatmap respectively for the two methods.

**Supplementary Figure 5.**

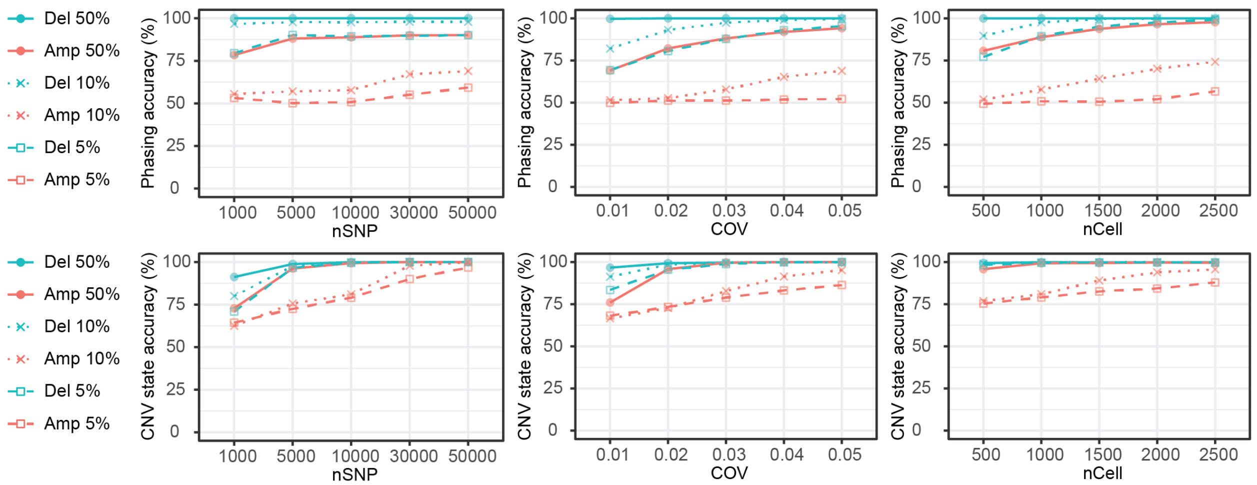

**Supplementary Fig. 5: Power for the detection of 1 copy deletion and 1 copy amplification for data of varying coverage (per base), heterozygous SNP count, and number of cells.** The heterozygous SNP count reflects the size of the region: larger regions contain more heterozygous loci. Cells were clustered based on the minimum distance of $\hat{\theta}_i$ to the canonical values. Top: phasing accuracy, defined as the proportion of SNPs with $\hat{I}_j$ correctly estimated; bottom: cell CNV state accuracy, defined as the proportion of cells that are correctly assigned to carrier state. Amp: amplification. Del: deletion. Line types represent different proportions (50%, 10% and 5%) of carrier cells. The number of SNPs, coverage, number of cells and purity were set as 10,000, 0.03, 1000, and 0.5 if not specified.

#### Supplementary Figure 6.

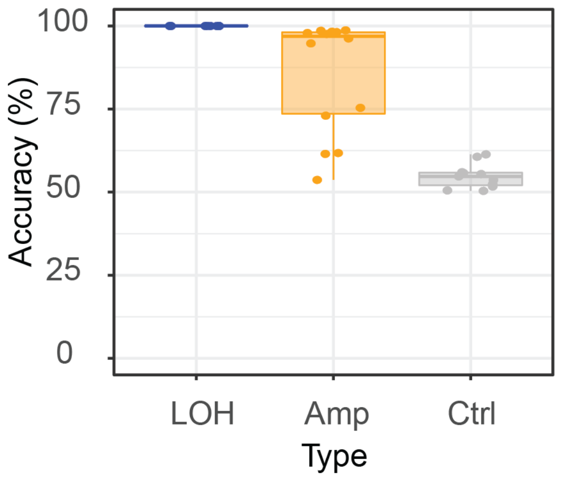

**Supplementary Fig. 6: Phasing accuracy for the CNA regions in the P6198 sample by comparing to the matched linked-read sequencing data.** LOH: segments with any LOH events. Amp: segments with amplifications that lead to allelic imbalance. Ctrl: control segments without allelic imbalance.

#### Supplementary Figure 7.

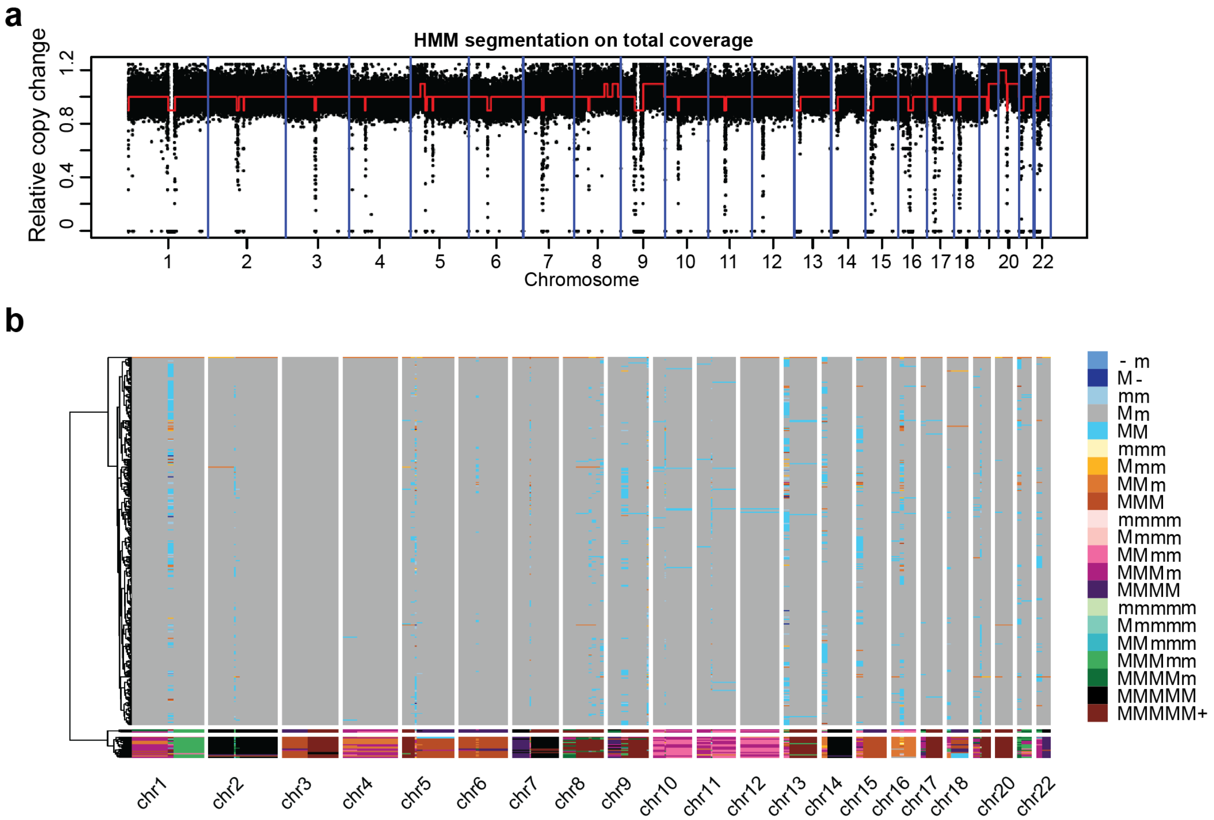

**Supplementary Fig. 7: The segmentation plot (a) and heatmap of the genome-wide haplotype profiles (b) for the P5846 sample.** In the color panel, M and m represent the “Major haplotype” and “minor haplotype” respectively.

**Supplementary Figure 8.**

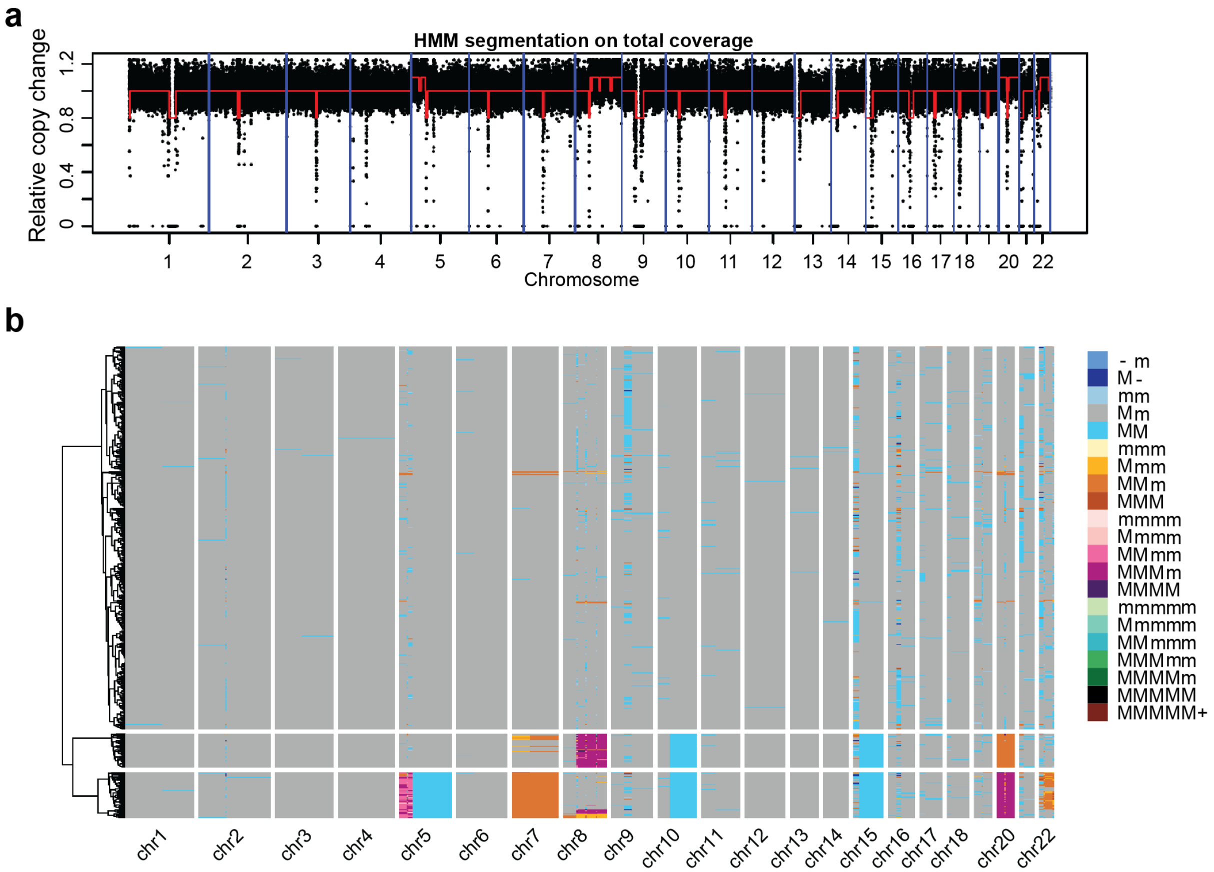

**Supplementary Fig. 8: The segmentation plot (a) and heatmap of the genome-wide haplotype profiles (b) for the P5847 sample.** In the color panel, M and m represent the “Major haplotype” and “minor haplotype” respectively.

#### Supplementary Figure 9.

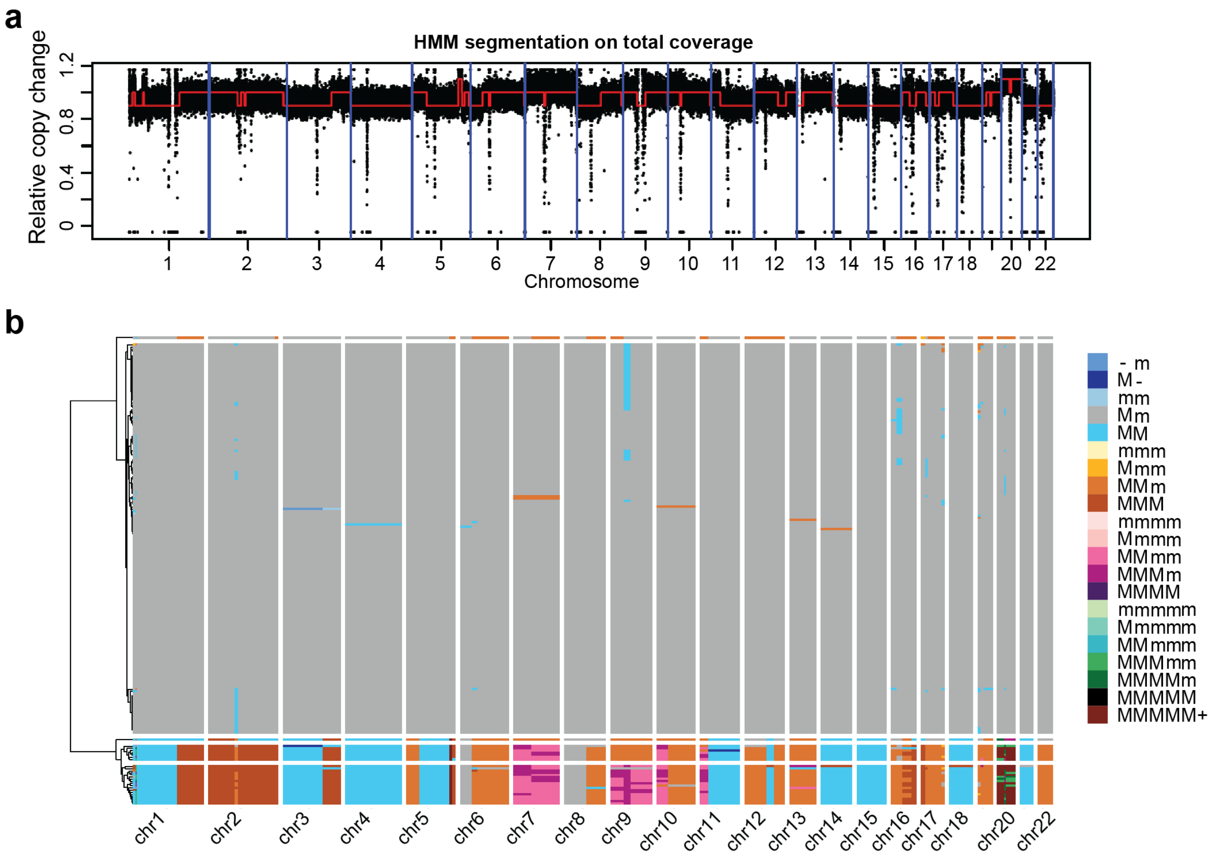

**Supplementary Fig. 9: The segmentation plot (a) and heatmap of the genome-wide haplotype profiles (b) for the P5915 sample.** In the color panel, M and m represent the “Major haplotype” and “minor haplotype” respectively.

#### Supplementary Figure 10.

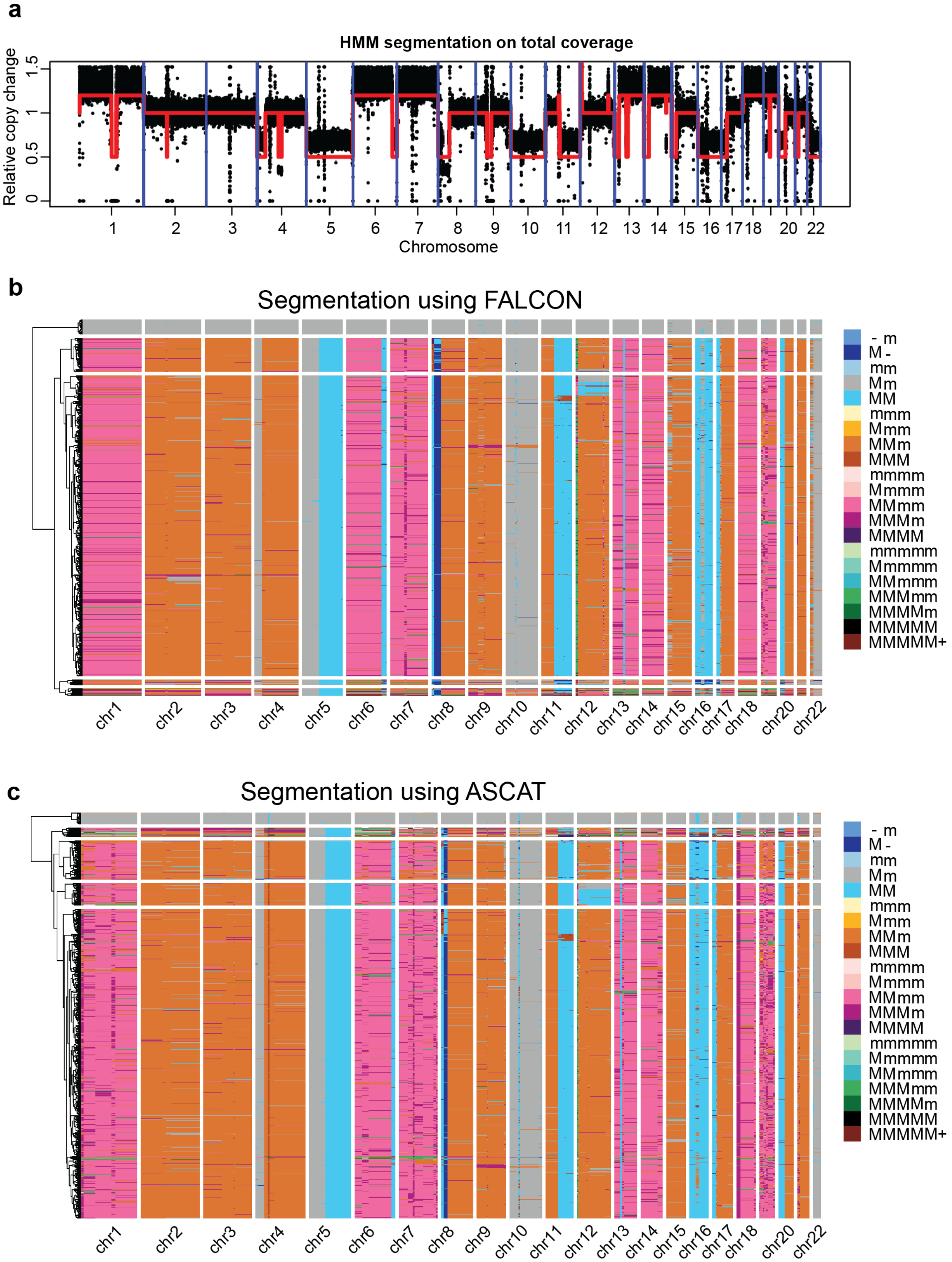

**Supplementary Fig. 10: The segmentation plot (a) and heatmap of the genome-wide haplotype profiles (b&c) for the P6198 sample.** Two segmentation methods were applied on this sample. (b) The genome-wide haplotype profiles estimated using FALCON. (c) The genome-wide haplotype profiles estimated using ASCAT. In the color panel, M and m represent the “Major haplotype” and “minor haplotype” respectively.

**Supplementary Figure 11.**

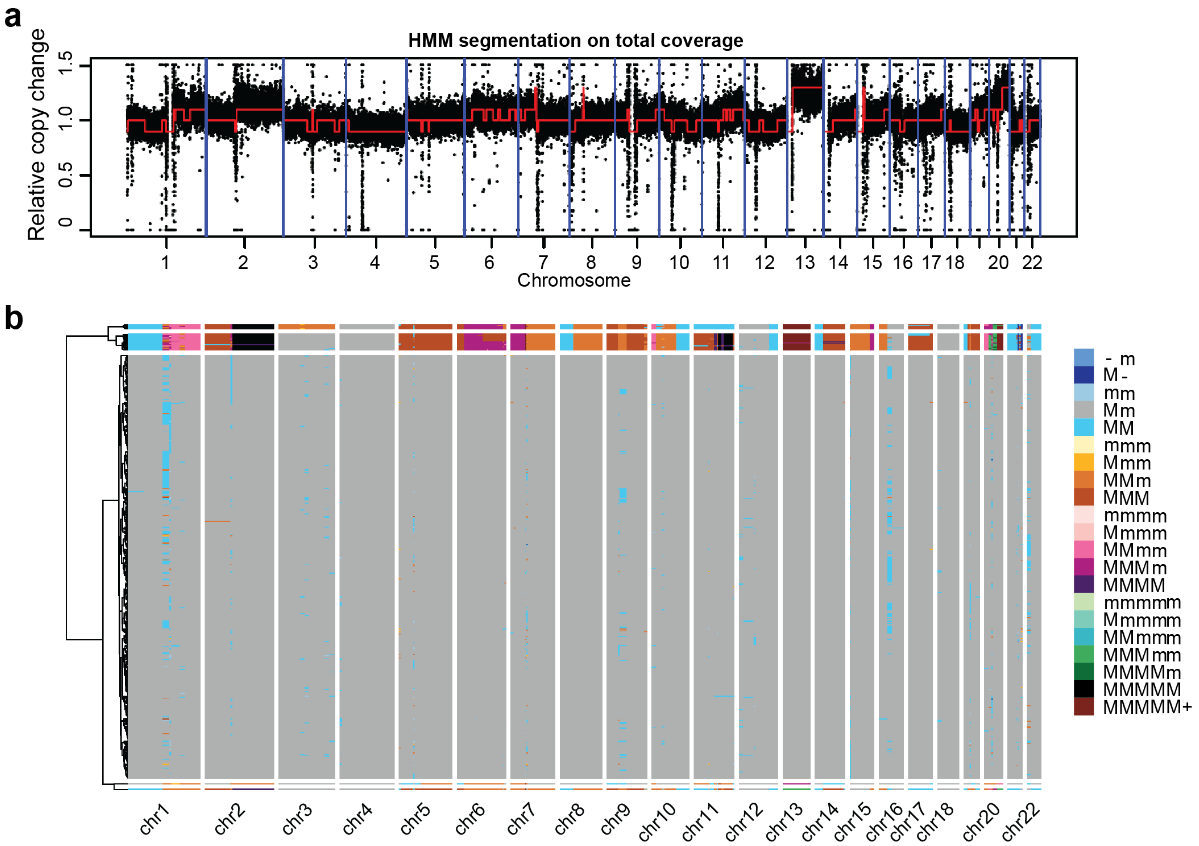

**Supplementary Fig. 11: The segmentation plot (a) and heatmap of the genome-wide haplotype profiles (b) for the P6335 sample.** In the color panel, M and m represent the "Major haplotype" and "minor haplotype" respectively.

#### Supplementary Figure 12.

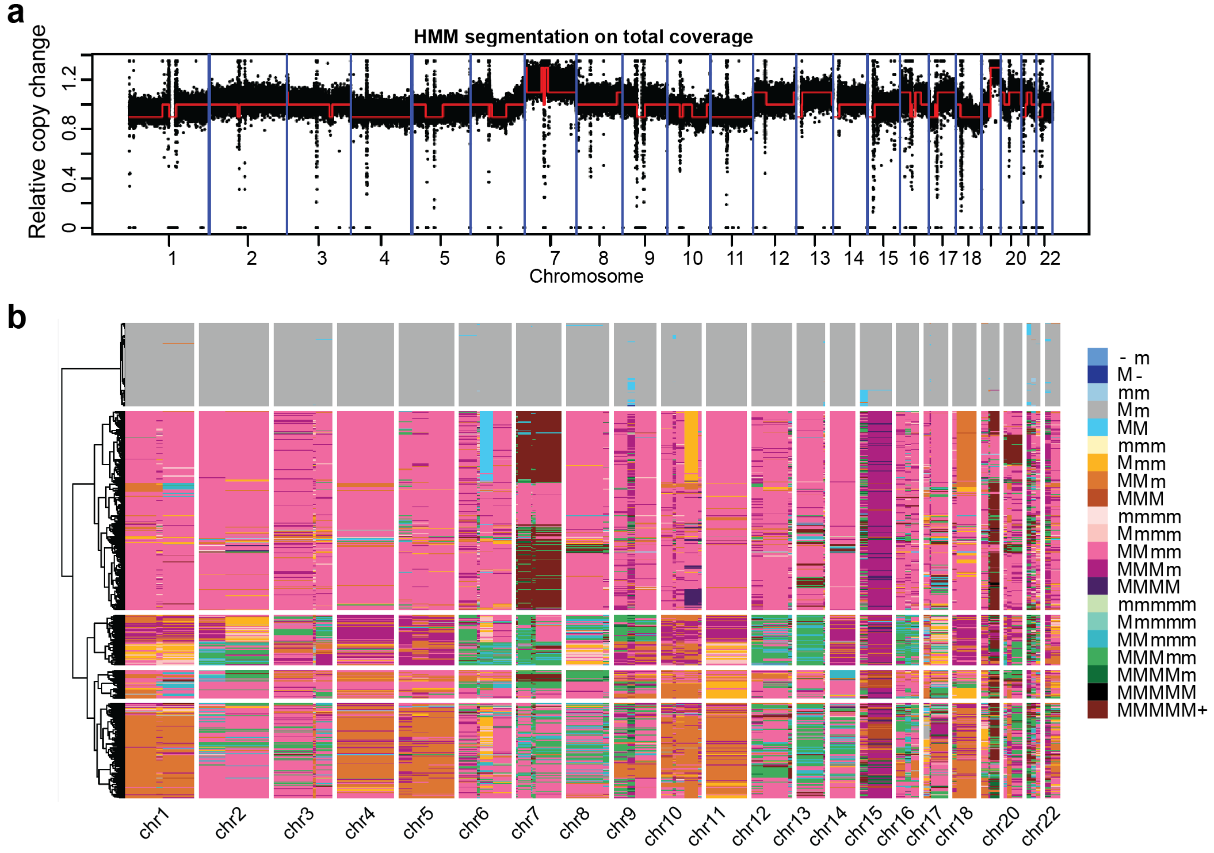

**Supplementary Fig. 12: The segmentation plot (a) and heatmap of the genome-wide haplotype profiles (b) for the P6461 sample.** In the color panel, M and m represent the “Major haplotype” and “minor haplotype” respectively.

#### Supplementary Figure 13.

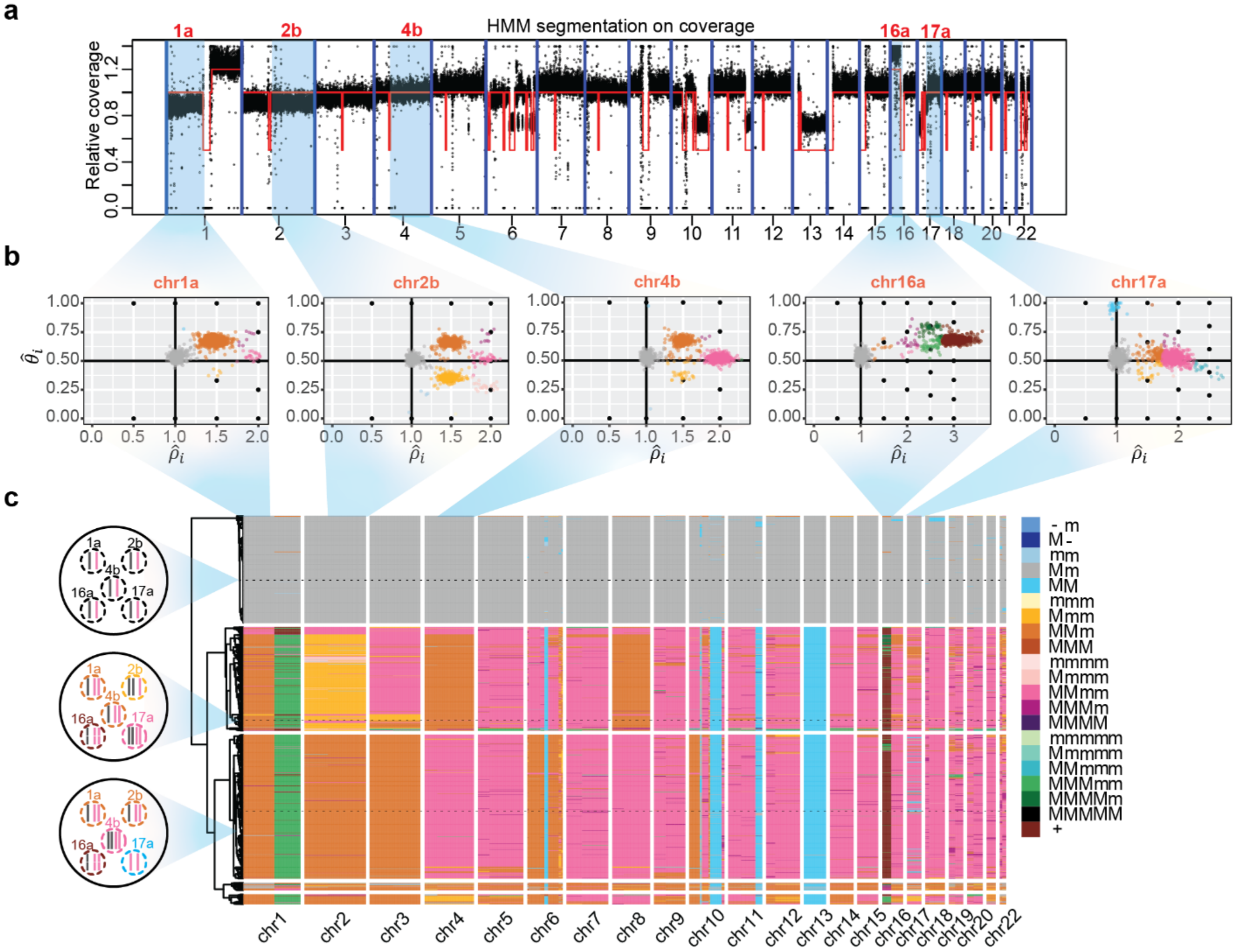

**Supplementary Fig. 13: The segmentation plot and heatmap of the genome-wide haplotype profiles for the breast cancer sample (section D).** (a) Genome segmentation using HMM on the pooled coverage signals across the cells. (b) Genotype profiles of five example regions. The coloring scheme is same as that in part (c). (c) Hierarchical clustering of single-cell ASCN genotypes reveals complex subclone structure. Genotypes of the five regions in three example cells from the three major subclones are shown in the left. Different colors represent different genotypes. In the color panel, M and m represent the “Major haplotype” and “minor haplotype” respectively.

**Supplementary Figure 14.**

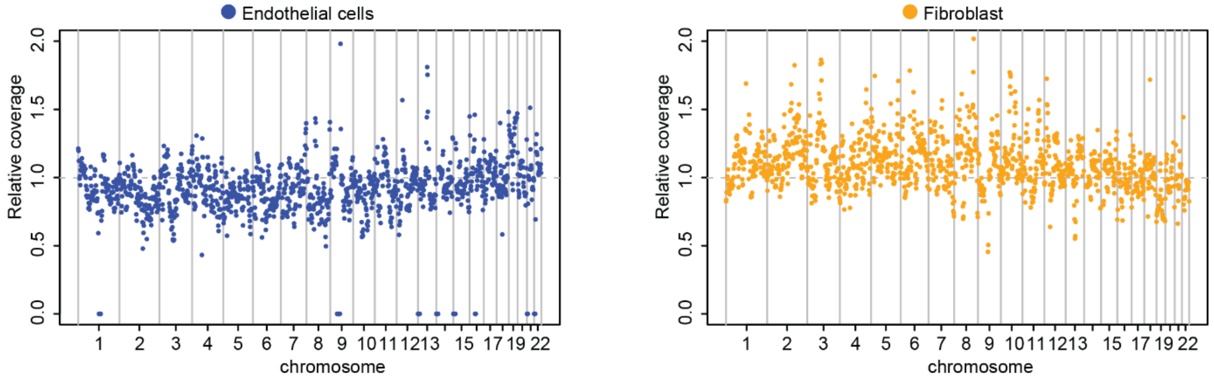

**Supplementary Fig. 14. Genome-wide coverage comparison in large genomic bins between two normal cell types for the SU008 sample.** Each point in the scatter plots represent the normalized read counts in each 10Mb bin along the genome for endothelial cells (left) and fibroblasts (right). The normalized read counts were computed by dividing the median read counts of cells of one normal cell type by the median read counts of cells of the other cell type.

**Supplementary Figure 15.**

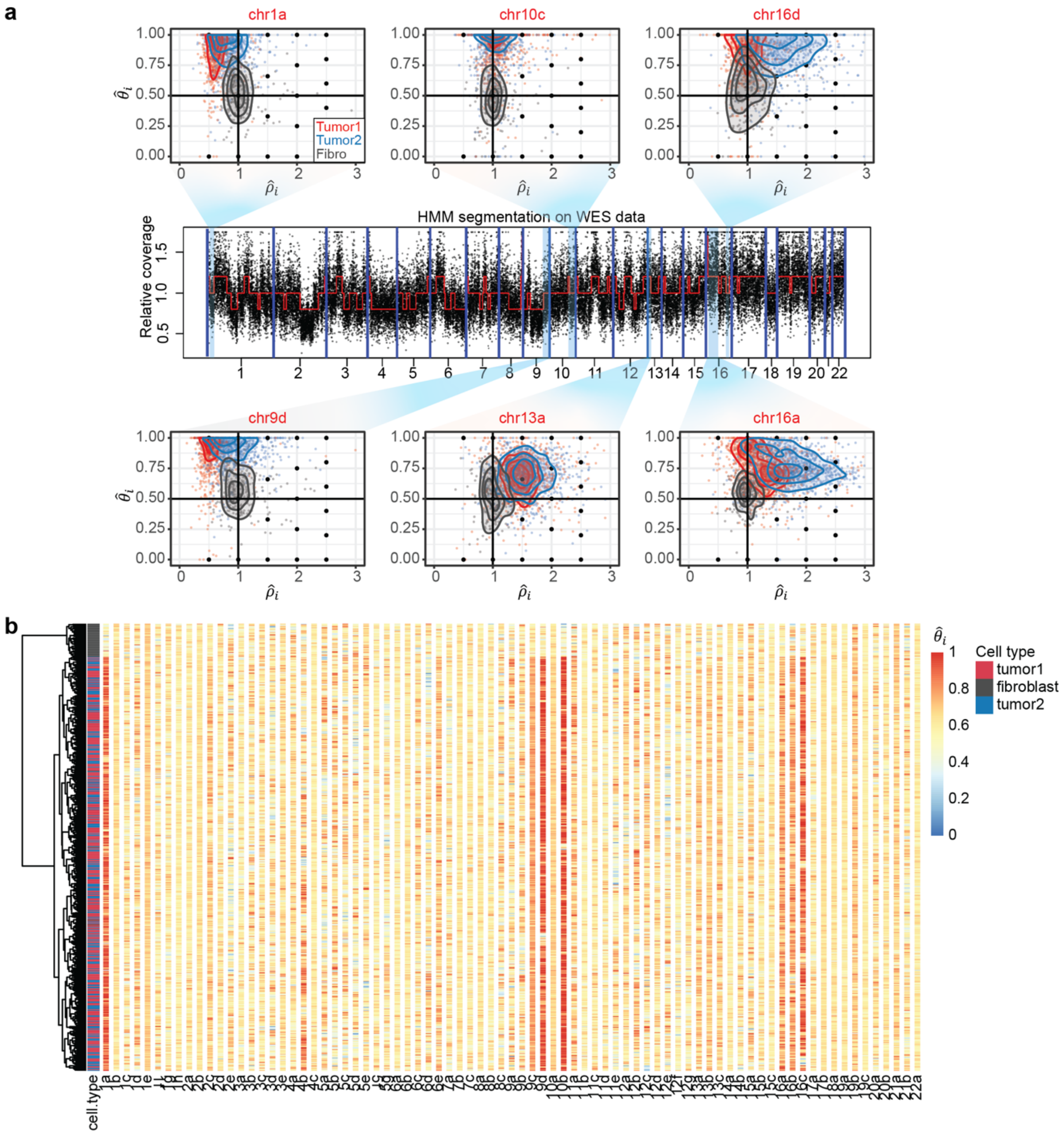

**Supplementary Fig. 15: Single cell genotyping of CNV events by Alleloscope for scATAC-seq data of a basal cell carcinoma sample (SU006[1]).** (a) Genotype profiles of six example regions. The regions were taken from the segmentation of whole exome sequencing (WES) data. Each dot represents a cell-specific $(\hat{\rho}_i,\ \hat{\theta}_i)$ pair. Cells are colored by annotation derived from peak signals[1]. Two tumor cell clusters, identified using ATAC peaks, are labeled by red and blue; fibroblasts (Fibro) are labeled by grey. Density contours of the three cell subpopulations are also shown. (b) Hierarchical clustering of cells in scATAC-seq by $\hat{\theta}_i$ reveals that the two tumor subpopulations are differentiated by peak signals that don't correlate with broad copy number events.

**Supplementary Figure 16.**

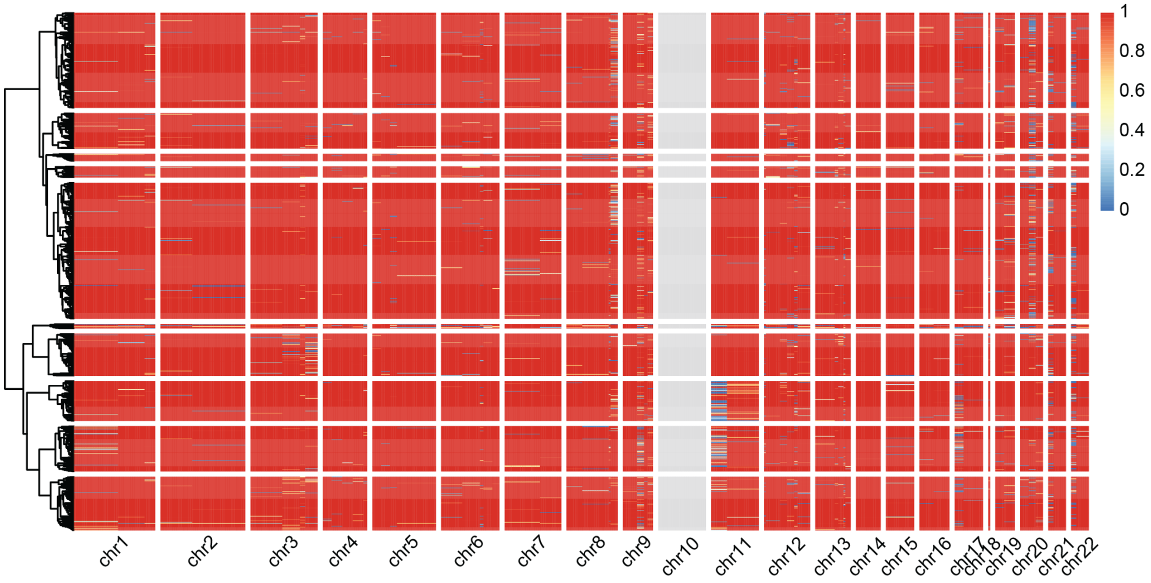

**Supplementary Fig. 16: Confidence scores for the genotype assignment of each cell in each region for the SNU601 scDNA-seq dataset.**

#### Supplementary Figure 17.

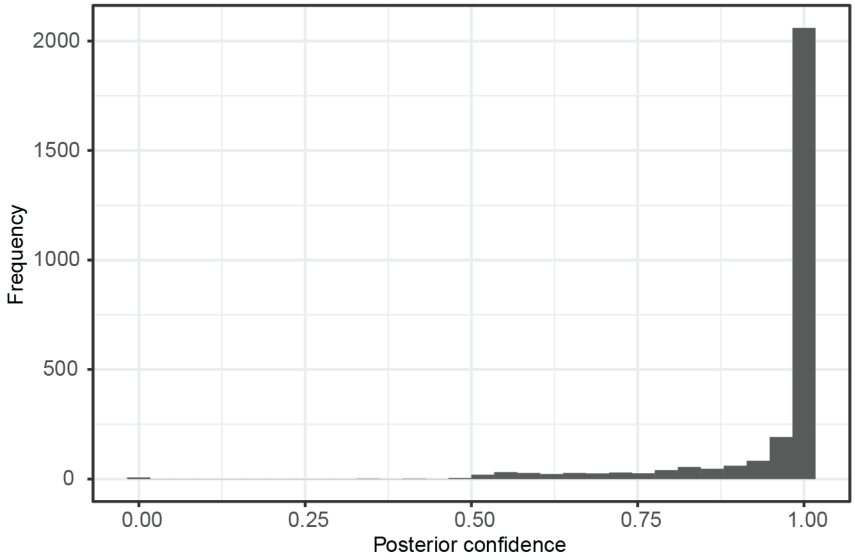

**Supplementary Fig. 17: Distribution of the posterior confidence scores of subclone assignment for the 2,753 cells from SNU601 scATAC-seq.**

**Supplementary Figure 18.**

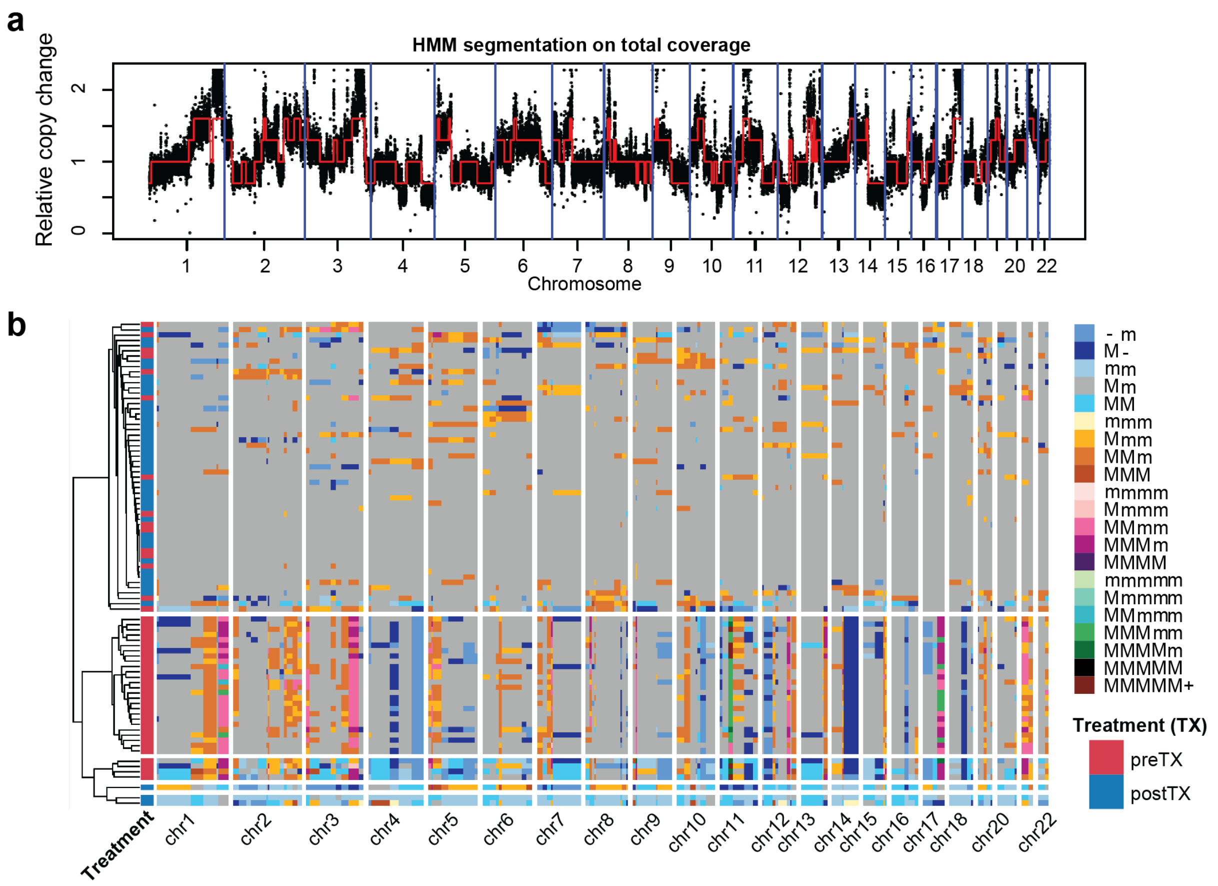

**Supplementary Fig. 18: The segmentation plot (a) and heatmap of the genome-wide haplotype profiles (b) for the HM-SNS sample[2].** In the color panel, M and m represent the “Major haplotype” and “minor haplotype” respectively.

#### Supplementary Tables

**Supplementary Table 1. Summaries of the scDNA-seq datasets.**

| Sample | Cancer type | Source | MSI status | Paired normal | Linked -reads | Coverage per cell | Cell number | Ref |
|---|---|---|---|---|---|---|---|---|
| P5846 | Gastric | Primary tissue | MSS | Yes | No | 454,806 | 510 | [3] |
| P5847 | Gastric | Primary tissue | - | Yes | No | 422,134 | 715 | - |
| P5915 | Colorectal | Liver meta | MSS | Yes | Yes | 1,262,629 | 233 | [3] |
| P5931 | Gastric | Primary tissue | MSI | Yes | Yes | 730,932 | 796 | - |
| P6198 | Colorectal | Liver meta | MSS | Yes | Yes | 532,343 | 2,271 | [3] |
| P6335 | Colorectal | Omentum meta | MSS | No | Yes | 564,058 | 953 | [3] |
| P6461 | Colorectal | Primary tissue | - | Yes | Yes | 483,524 | 1,242 | - |
| SNU601 | Gastric | Ascites meta | MSS | No | No | 565,648 | 1,531 | [4] |
| BC10x (secD) | Breast | Primary tissue | - | No | No | 781,506 | 1,916 | [5] |
| BC10x (secE) | Breast | Primary tissue | - | No | No | 951,225 | 2,053 | [5] |

**Supplementary Table 2. Performance of CHISEL with or without external phasing information.** CHISEL was run with default setting (50kb block size with external phasing information) or without external phasing (block size 1)

| Dataset | Method | Sensitivity | Specificity |
|---|---|---|---|
| P5931 | CHISEL: default (before correction) | 0.7520 | 0.9434 |
| | CHISEL: default (after correction) | 0.0112 | 0.9508 |
| | CHISEL: block size 1 (before correction) | 0.5939 | 0.7878 |

| | | | |
|---|---|---|---|
| | CHISEL: block size 1 (after correction) | 0.7465 | 0.7948 |
| P6198 | CHISEL: default (before correction) | 0.9397 | 0.9311 |
| | CHISEL: default (after correction) | 0.9700 | 0.9359 |
| | CHISEL: block size 1 (before correction) | 0.8067 | 0.8608 |
| | CHISEL: block size 1 (after correction) | 0.8320 | 0.8686 |
| P6335 | CHISEL: default (before correction) | 0.7858 | 0.9873 |
| | CHISEL: default (after correction) | 0.8404 | 0.9943 |
| | CHISEL: block size 1 (before correction) | 0.6959 | 0.9705 |
| | CHISEL: block size 1 (after correction) | 0.7450 | 0.9782 |

**Supplementary Table 4. Summaries of the scATAC-seq datasets.**

| Sample | Cancer type | Source | Matched DNA | Coverage per cell | Cell number | Ref |
|---|---|---|---|---|---|---|
| SU006 | Basal cell carcinoma | Primary tissue | Bulk WES | 41,368 | 2771 | [1] |
| SU008 | Basal cell carcinoma | Primary tissue | Bulk WES | 36,057 | 788 | [1] |
| SNU601 | Gastric | Ascites meta | scDNA-seq | 73,845 | 3614 | - |

#### Supplementary Results

##### Benchmark 1: Assessment of Alleloscope and CHISEL using whole genome haplotypes derived from linked read sequencing

So far, CHISEL is the only other comparable method for allele-specific copy number estimation with scDNA-seq data. We comprehensively benchmarked Alleloscope and CHISEL using the following two strategies: (1) We compared each method's output under default parameter settings, without linked read phasing, to the same method's output obtained given linked-reads phasing. This was done for the five samples (P5931, P5915, P6198, P6335, and P6461) of varying complexity for which matched linked-read sequencing was performed. (2) We compared each method's output at original sequencing coverage to the same method's output at 50% or 25% coverage. This was performed for the high-coverage breast cancer sample that CHISEL analyzed in their paper. The rationale is that results at original coverage, though noisy, should be closer to the truth than results at reduced coverage, and thus the method whose output remains more stable under data down-sampling is more accurate at these lower coverages (Supplementary Methods).

For the first benchmark strategy, we used P5931 as an illustrating example. P5931 tumor sample carries simpler but representative haplotype-specific profiles. For each method, we obtain two different sets of results: Results obtained by default parameter settings, and results obtained by plugging in known phases provided by the matched linked-read sequencing data. The latter, results obtained using known phases, is used as the gold standard for assessing each method. For CHISEL, there is an extra "correction step" where the inferred clones are used to generate consensus allele-specific copy number profiles for all cells within each clone.

Supplementary Fig. 2 compares the estimated genome-wide haplotype profiles by Alleloscope with and without known phases. As explained in the manuscript, Alleloscope's color scheme gives a different color for each allele-specific copy number state, with gray being normal diploid ("Mm"), gradients of brown being amplification, and gradients of blue being deletion. Alleloscope achieved a sensitivity of 0.9402 and specificity of 0.9986. The sensitivity here can also be referred to as "recall."

For CHISEL, there is an extra "correction step" where the inferred clones are used to generate consensus allele-specific copy number profiles for all cells within each clone. In Supplementary Fig 3, we show CHISEL's output with and without this correction step. For each result, CHISEL gives two heatmaps: the left heatmap showing the allele-specific copy number state, and the right heatmap showing, for configurations where there is an allelic imbalance, which is the major allele. Sensitivity for CHISEL is 0.7520 (before correction) and 0.0112 (after correction), and specificity is 0.9434 (before correction) and 0.9508 (after correction). Note that for this example, CHISEL's clone-based correction step actually erases most of the signal (the amplifications on chr7, chr8, and chr20, as well as the deletion on chr21 are erased by the correction). However, in other samples, we have noticed the reverse, where the results after the correction are better. Thus, CHISEL's estimates are not robust to errors in clonal inference. The better of CHISEL's two outputs (the uncorrected version) still has significantly lower specificity and sensitivity as compared to Alleloscope. As a sanity check, we also compared the two "gold standards": CHISEL's results obtained using linked-reads phasing, and Alleloscope's results obtained using linked-reads phasing. The two gold standards have a similarity of 0.9945 (before correction) and 0.9851 (after correction), thus indicating that they are both accurate reflections of the underlying truth.

Comparing Supplementary Figures 2 and 3, we can see how the results from Alleloscope and CHISEL differ. The cells in both plots are ordered by their ordering in the linked-read-based heatmap. In CHISEL's results, the cells deemed noisy are labeled with grey color in the clonal assignment for each result. There are many more "horizontal stripes", showing cells that have copy number estimates that disagree with its clonal average. rong clonal inference Also, there are many more cells deemed "too noisy" by CHISEL, these are clustered at the tops and bottoms of the heatmaps. This leads to the lowered specificity and sensitivity of CHISEL in this sample.

Besides using the default setting, we also explored the extent to which CHISEL relies on the external phasing information. We found that CHISEL is very sensitive to not only the block-size parameter, but also the selection of the phasing panel using P5931 as an example (Supplementary Fig. 3; Supplementary Table 2). Even for the same patient, the tumor tissue and matched normal tissue differ in their phasing profiles at key regions surrounding chromosomal breakpoints, which can lead to different results under CHISEL. Effects of external phasing were also explored in two additional samples—P6335 and P6198 (Supplementary Table 2). This reinforces the rationale that, for allele-specific copy number estimation of tumor samples, it would be best to not rely on external phasing.

We also performed such detailed assessment between CHISEL and Alleloscope on four other samples, with differing CNA complexity, that have matched linked-read sequencing data—P6198, P6335, P5915 and P6461. Sensitivity and specificity of the four samples and those of P5931 for the two methods are shown in Table 1. The results show that Alleloscope outperforms CHISEL in allele-specific copy number estimation for most samples, and both methods perform similarly for P6198. The high 0.97 sensitivity of CHISEL after correction results from the fact that P6198 is a tumor sample with one major subclone and multiple LOH regions that are easier to be detected (Supplementary Fig. 10). For P6198, CHISEL generates consensus allele-specific copy number profiles for all cells clustered in the same clone, which leads to extremely high sensitivity in this easier case.

**Benchmark 2: Assessment of method robustness by downsampling**

In addition to the linked-reads based benchmark, we also compared Alleloscope and CHISEL on the breast cancer sample that CHISEL analyzed in their study. This data set does not have true phasing information, but was sequenced at much higher coverage, and thus, we subsampled 50% and 25 % of the original dataset and compared the estimated results of the subsampled datasets to that of the original datasets to assess the performance of the two methods across varying sequencing coverage. The results on the original dataset, on the 50% subsampled dataset and on the 25% subsampled dataset, given by Alleloscope and CHISEL, are shown in Supplementary Fig. 4. For CHISEL, the clone-corrected output plots are shown since they are more accurate than the uncorrected ones. Using the results from the original dataset as the ground-truth, performance of Alleloscope and CHISEL for the two subsampled datasets is shown in Table 1. Sensitivity and specificity are both high (>0.90) for both Alleloscope and CHISEL on the 50% subsampled dataset. However, for the 25% subsampled dataset, sensitivity decreases to ~0.82 and specificity decreases to ~0.95 for Alleloscope, while CHISEL fails in estimating allele-specific copy numbers on this dataset with both sensitivity and specificity decreasing to ~65%. The performance can also be visualized in Supplementary Fig. 4. While the lineage plots look noisier with more dark horizontal bands as the coverage decreases, CHISEL fails to not only estimate allele-specific copy numbers for some regions (like those grey vertical bands on chr10, 11, 13, and 17), but also the direction of the two haplotypes for almost the whole genome.

For scDNA-seq allele-specific copy number estimation, Alleloscope has higher sensitivity and specificity comparing to the known phases provided by the matched linked-reads on the five scDNA-seq samples. The subsampling results also suggest that Alleloscope is more robust at lower coverage. **Algorithmic differences between CHISEL and Alleloscope**

Here we describe the algorithmic differences between CHISEL and Alleloscope, which can help explain the differences in their performance. CHISEL was designed for scDNA-seq data and is not applicable to scATAC-seq data. The improvements of Alleloscope over CHISEL are due to fundamental differences in their algorithm design: Alleloscope is a top-down method that first segments the genome, allowing for the aggregation across all SNPs in a large region to estimate haplotype profiles for each cell. This allows the algorithm to achieve higher sensitivity for events carried by smaller subclones, as well as robustness against local fluctuations in allelic coverage. Delineating the segments in the first step also allows for the simultaneous estimation of phase and single cell haplotype ratios, thus bypassing the need for external phasing data. In comparison, CHISEL is a bottom-up approach that estimates copy number profiles of each cell in smaller fixedlength bins (default 5Mb). Selection of the bin size faces a trade-off between having more SNPs (and thus more data points for estimation) and heterogeneity of the true CNA profile in the within the bin. To overcome the sparsity of data within each bin, CHISEL relies on external phasing to achieve better allele-specific copy number estimation. Choice of the external phasing data can substantially impact estimation results (Supplementary Fig. 3).

#### Supplementary Methods

##### Second-stage estimation

To improve copy number state estimation in regions with low proportion of cells carrying CNAs, a second-stage scheme was developed. We first estimated cell-level CNV states by the methods described in Methods. After the first-round estimation, some cells with low proportions might have shifts in their coverage ($\hat{\rho}_{ir}$) but their allelic imbalance level ($\hat{\theta}_{ir}$) do not correspond to the coverage change for some regions. Visualizing this from the scatter plots, the second-stage estimation can be executed to estimate phases using only the targeted cells.

In the second-round estimation, SNPs in region $r$ are first filtered out if no read is observed among the targeted cells. The phases of these filtered SNPs can be estimated as described in the "*SNP Phasing and Single-cell Allele Profile Estimation per region"* section of Methods. Using the estimated phases for the filtered SNPs, the allelic imbalance level of the targeted cells and other non-targeted cells can both be estimated.

##### Benchmarks of Alleloscope and CHISEL

We benchmarked Alleloscope and CHISEL on the four scDNA-seq samples using two strategies explained below. These samples include five (P5931, P5915, P6198, P6335, and P6461) for which matched linked-read sequencing was performed, and the high-coverage breast cancer sample that CHISEL analyzed in their paper.

For CHISEL, we prepared the four required input files following the online tutorial. Since no matched normal sample exits for P6335 and BC10x, we first generated bam files for normal cells in the tumor samples using CHISEL as they suggested. To run CHISEL with the default settings, Eagle2 through Michigan Imputation Server was used to phase germline SNPs following their pipeline.

*Benchmark 1: Comparison to results obtained by linked-reads phase information.*

For each method, we obtain two different sets of results: Results obtained by default parameter settings, and results obtained by plugging in known phases provided by the matched linked-read sequencing data. The latter, results obtained using known phases, is used as the gold standard for assessing each method. Since linked-read sequencing data can provide phase information covering Mb scale, haplotype block was set as 1Mb for P5931 to run CHISEL to retrieve the result used as the gold standard. For P6198 and P6335, we observed that setting a block size of 1Mb introduced additional noise because the 1Mb block used might be different from that provided by the linked-read sequencing data. We instead used the original 50kb block for these two samples. GRCh38 referecne genome was used to analyze these samples. Between the results by default setting and by plugging in known phases, the directions of the two haplotypes (considered as either A or B) might be reversed for some chromosomes depending on the phasing information used. To make the results comparable, directions of the two haplotypes estimated by the default setting were used as the scaffold to place the directions of the two haplotypes in the results by plugging the known phases, which was similar to the comparison in Alleloscope.

*Benchmark 2: Comparison of results at down-sampled coverage to results at original coverage for a high coverage sample.*

We compared Alleloscope and CHISEL on the breast cancer sample that CHISEL analyzed in their study by subsampling. We first subsampled 50% and 25 % of the original dataset using samtools. Alleloscope and CHISEL were run on the original dataset, 50% and 25% subsampled datasets using default settings on the GRCh37 reference genome. The estimated results of the subsampled datasets were compared to that of the original datasets (used as the gold standard) to assess the performance of the two methods across varying sequencing coverage respectively.

To compare Alleloscope and CHISEL, each segment analyzed by Alleloscope was divided into 5Mb bins following the format in CHISEL's output. Then, sensitivity and specificity were computed as follows:

$$Sensitivity = \frac{TP}{P},$$

where:

TP 5Mb bins across the cells and across the genome considered to be abnormal (haplotype profiles other than 1|1) in both the results used as gold standard and the estimated results;

P: 5Mb bins across the cells and across the genome considered to be abnormal in the result used as gold standard.

and

$$Specificity = TN/N,$$

where

TN: 5Mb bins across the cells and across the genome considered to be normal diploid in both the results used as gold standard and the estimated results;

N: 5Mb bins across the cells and across the genome considered to be normal diploid in the result used as gold standard.

For a sanity check, similarity was also computed for the two “gold standards”: CHISEL’s results obtained using linked-reads phasing, and Alleloscope’s results obtained using linked-reads phasing. We first generated two cell by 500M-bin matrices for both methods with the values indicating the allele-specific copy number profiles. Then similarity was computed as the proportion of the same copy number profiles across the cells and the 500M bins.

**Assessment of coverage in large genomic bins for scATAC-seq**

To compare with the CNA analysis method using only coverage for the scATAC-seq data, we assessed if false-positive signals can still be observed even with large genomic bins using the two normal cell types (endothelial cells and fibrobalsts) in the SU008 scATAC-seq dataset. The identity of the two normal cell types were based on their genome-wide peak signals. Following the method in the original paper[1], we first summed all the read signals normalized with the cell size in each 10Mb bins with a 2Mb sliding window along the genome for each cell. Then for each nor mal cell type, the signals of each bin were normalized by dividing the median of the total signals across the cells in one normal cell type by the median of the total signals across the cells in the other cell type. This step was similar to using normal cells as the control to normalize the signals in tumor cells to assess the copy number change.

**Inference of MLE estimates for the copy number adjustment model**

For peak *k*, the binomial distribution with specific terms to adjust for copy numbers is used to model the observed read counts:

$$Y_{ck} \sim Binomial(N_c,\ f_k(\theta_c)p_{ck}),$$

see details of each term in Methods. For each peak $k$, two clones (clone $c_1$ and clone $c_2$) are compared using the generalized likelihood ratio test (GLRT) with the hypothesis $H_0: p_{c_1k} = p_{c_2k} = p_0$ and $H_1: p_{c_1k} \neq p_{c_2k}$. as the GLRT in this case has null distribution that is $\chi_1^2$, and has the form:

$$LLR_k = 2\ell(\hat{p}_{c_1k}, \hat{p}_{c_2k}) - 2\ell(\hat{p}_{0k})$$

Where $2\ell(\hat{p}_{c_1k}, \hat{p}_{c_2k})$ is the maximized log-likelihood under the alternative, with

$$\hat{p}_{c_1,k} = \frac{Y_{c_2k}}{N_{c_2}}, \qquad \hat{p}_{c_1,k} = \frac{Y_{c_2k}}{N_{c_2}}$$

and $l(\hat{p}_{0k})$ is the maximized log-likelihood under the null, with the MLE computed as follows: For simplicity, we first use $p_0$ to denote $p_{0k}$; $f_1, N_1, Y_1$ to denote $f_k(\theta_{c_1}), N_{c_1k}, Y_{c_1k}$; and $f_2, N_2, Y_2$ to denote $f_k(\theta_{c_2}), N_{c_2k}, Y_{c_2k}$. Then

$$\ell(p_0) = \log[(f_1p_0)^{Y_1}(1 - f_1p_0)^{N_1-Y_1}(f_2p_0)^{Y_2}(1 - f_2p_0)^{N_2-Y_2}]$$

$$\frac{d}{d(p_0)}\ \ell(p_0) = \frac{Y_1f_1}{f_1p_0} + (N_1 - Y_1)\left(-\frac{f_1}{1 - f_1p_0}\right) + \frac{Y_2f_2}{f_2p_0} + (N_2 - Y_2)\left(-\frac{f_2}{1 - f_2p_0}\right)$$

$$= \frac{Y_1}{p_0} - (N_1 - Y_1)\frac{f_1}{1 - f_1 p_0} + \frac{Y_2}{p_0} - (N_2 - Y_2)\frac{f_2}{1 - f_2 p_0} = 0$$

$$Y_1(1 - f_1 p_0)(1 - f_2 p_0) - p_0 f_1 (N_1 - Y_1)(1 - f_2 p_0) + Y_2(1 - f_1 p_0)(1 - f_2 p_0)$$
$$- p_0 f_2 (N_2 - Y_2)(1 - f_1 p_0) = 0,$$

$$[Y_1 f_1 f_2 {p_0}^2 - Y_1(f_1 + f_2)p_0 + Y_1] - p_0 f_1 N_1 + p_0 f_1 N_1 f_2 p_0 + p_0 f_1 Y_1 - p_0 f_1 Y_1 f_2 p + [Y_2 f_1 f_2 {p_0}^2$$
$$- Y_2(f_1 + f_2)p_0 + Y_2] - p_0 f_2 N_2 + p_0 f_2 N_2 f_1 p_0 + p_0 f_2 Y_2 - p_0 f_1 Y_2 f_2 p = 0,$$

$$(Y_1 + Y_2) f_1 f_2 {p_0}^2 - (Y_1 + Y_2) p_0 (f_1 + f_2) + (Y_1 + Y_2) - p_0 f_1 (N_1 - Y_1) - p_0 f_2 (N_2 - Y_2)$$
$$+ {p_0}^2 f_1 f_2 (N_1 - Y_1) + p_0 f_2 (N_2 - Y_2)$$
$$= f_1 f_2 (N_1 + N_2) {p_0}^2 - (f_1 N_1 + f_2 N_2 + f_1 Y_2 + f_2 Y_1) p_0 + (Y_1 + Y_2) = 0,$$

Let $a = f_1 f_2 (N_1 + N_2); b = (f_1 N_1 + f_2 N_2 + f_1 Y_2 + f_2 Y_1); c = (Y_1 + Y_2)$,

$$\hat{p}_0 = \frac{-b - \sqrt{b^2 - 4ac}}{2a}, \left(0 < p_0 < min\left\{\frac{1}{f_1}, \frac{1}{f_2}\right\}\right).$$

**Highly-multiplexed single-nucleus-sequencing (HM-SNS) data preprocessing and analysis**

To show that Alleloscope can work on scDNA-seq data generated using protocols other than the 10x one, we applied the method to a HM-SNS dataset form Kim et al.'s study. Total 89 fastq files for patient 2 were retried from SRP114962. Raw fastq files were aligned to the GRCh37 reference genome using bwa-mem with duplicate reads removed using the Picard toolkits. SAMtools was used to add read group, sort and index the bam files. After the bam files for individual cells were generated, the standard GATK workflow was followed to joint-call SNPs for all the cells to generate the SNP by cell matrices for the reference and alternative alleles. The bin by cell matrix was generated using SCOPE[6]. To obtain a SNP set including only SNVs that are more possible to be germline SNPs, we further filtered out the SNVs <10 reads or with extreme VAF values outside the (0.3, 0.9) range. A 0.3 threshold was due to a small peak in the histogram of the VAF values below 0.3, which resulted from many somatic SNVs detected when calling SNVs on individual cells. Since no matched normal exists for this sample, we first identified tumor and normal cells using $\hat{\theta}_{ir}$ estimated using each chromosome (instead of using the segments) in the sample for the segmentation step.
